# Structural basis for the transport mechanism of the human glutamine transporter SLC1A5 (ASCT2)

**DOI:** 10.1101/622563

**Authors:** Xiaodi Yu, Olga Plotnikova, Paul D. Bonin, Timothy A. Subashi, Thomas J. McLellan, Darren Dumlao, Ye Che, Yinyao Dong, Elisabeth P. Carpenter, Graham M. West, Xiayang Qiu, Jeffrey S. Culp, Seungil Han

## Abstract

Alanine-serine-cysteine transporter 2 (ASCT2, SLC1A5) is the primary transporter of glutamine in cancer cells and regulates the mTORC1 signaling pathway. The SLC1A5 function involves finely tuned orchestration of two domain movements that include the substrate-binding transport domain and the scaffold domain. Here, we present cryo-EM structures of human SLC1A5 and its complex with the substrate, L-glutamine in an outward-facing conformation. These structures reveal insights into the conformation of the critical ECL2a loop which connects the two domains, thus allowing rigid body movement of the transport domain throughout the transport cycle. Furthermore, the structures provide new insights into substrate recognition, which involves conformational changes in the HP2 loop. A putative cholesterol binding site was observed near the domain interface in the outward-facing state. Comparison with the previously determined inward-facing structure of SCL1A5 provides a basis for a more integrated understanding of substrate recognition and transport mechanism in the SLC1 family. Our structures are likely to aid the development of potent and selective SLC1A5 inhibitors for the treatment of cancer and autoimmune disorders.

## Introduction

SLC1A5 (ASCT2) catalyzes an obligatory Na^+^-dependent antiport in which Na^+^ together with an extracellular neutral amino acid are exchanged with an intracellular neutral amino acid (1, 2). It is the only glutamine transporter of the solute carrier family 1 (SLC1) family and functions as the primary transporter of glutamine in cancer cells (3). Inhibition of SLC1A5 glutamine transport by small molecules and shRNA-mediated SLC1A5 knockdown have been shown to decrease endometrial cancer cell growth (4). Similarly, SLC1A5 inhibition and knockdown suppressed mTORC1 signaling and induced rapid cell death in breast cancer cells (5). In addition, SLC1A5 ^−/−^ mice are deficient in T cell receptor-mediated activation of the metabolic kinase mTORC1 and as a result exhibit improved clinical scores in experimental autoimmune animal models of multiple sclerosis and colitis (6).

The structure and function of SLC1 transporters have been studied using bacterial aspartate transporter homologues Glt_PH_ from *Pyrococcus horikoshii* (7–12), GltTK from *Thermococcus kodakarensis* (13, 14), human glutamate transporter SLC1A3 (15) and glutamine transporter SLC1A5 (16). All of these transporters form a trimer of independently functioning protomers that uses an elevator mechanism to carry amino acids across membranes (11, 17). Each protomer has a scaffold domain and a movable transport domain which slides and carries the solute. The substrate transport is initiated by its binding, followed by the conformational transitions of the transporter between an outward- and inward-facing states with the substrate binding site accessible to the extracellular surface or to the intracellular surface, respectively (11, 17). Glt_PH_ structures have been reported in both the outward- and the inward-facing states (8, 9, 11, 18, 19). Crystal structures of unbound and substrate-bound Glt_TK_ have been reported in the outward-facing conformation. The crystal structures of SLC1A3 were determined with competitive and allosteric inhibitors in the outward-facing conformation (15). The technical challenges for crystallographic studies of this gene family are clearly evident from the reports on the SLC1A3 crystal structures which required mutation of nearly 25% of the native amino acids to create a thermostable protein for crystallization. Recently, the cryo-EM structure of SLC1A5 in complex with glutamine was reported in the inward-facing state (16). In order to gain insights into structural features that permit substrate recognition in outward-facing state, using an antibody fragment as a fiducial marker we have solved the cryo-EM structures of the unliganded transporter and its complex with glutamine at 3.5 and 3.8 Å resolution, respectively. Herein we describe key structural elements that facilitate the transition between inward- and outward-facing states and the identification of a putative allosteric binding pocket of SLC1A5. Additional biochemical, biophysical and structural studies provide insights for substrate specificity may also aid the development of potent and selective SLC1A5 inhibitors for treating cancer and autoimmune disorders.

## Results

### Function and overall architecture

Our full-length SLC1A5 construct showed high expression, stability and transport function (SI Appendix, Fig. S1a-d). The presence of affinity tags used for purification did not affect the sodium-dependent glutamine uptake when HAP1 SLC1A5 knock-out cells were transiently transfected with full-length SLC1A5 (Fig. 1a). To capture an outward-facing state of the transporter, we generated its complex with a commercially available cKM4012 Fab fragment. Negative-stain electron microscopy revealed that one molecule of Fab was bound to each protomer of a SLC1A5 homotrimer (SI Appendix, Fig. S1e-f). The three-fold symmetry was broken due to the dynamic nature of each Fab and protomer. Single-particle electron cryo-microscopy (cryo-EM) analysis without imposing any symmetry restraint resulted in a facile three-dimensional reconstruction at an overall resolution of 3.5 Å (Fourier shell correlation (FSC) = 0.143 criterion; SI Appendix, Fig. S2). The highly flexible Fab fragments were masked out to maximize the structural quality of SLC1A5 (SI Appendix, Fig. S2-3). The quality of the EM density map was sufficient to allow model building for residues 43-488 (SI Appendix, Fig. S4, Table S1). The limited resolution on the Fab region prevents a detailed and accurate analysis of the protein-fab interface. The primary purpose of the Fab was to serve as a fiducial marker for particle alignment.

**Figure 1.**
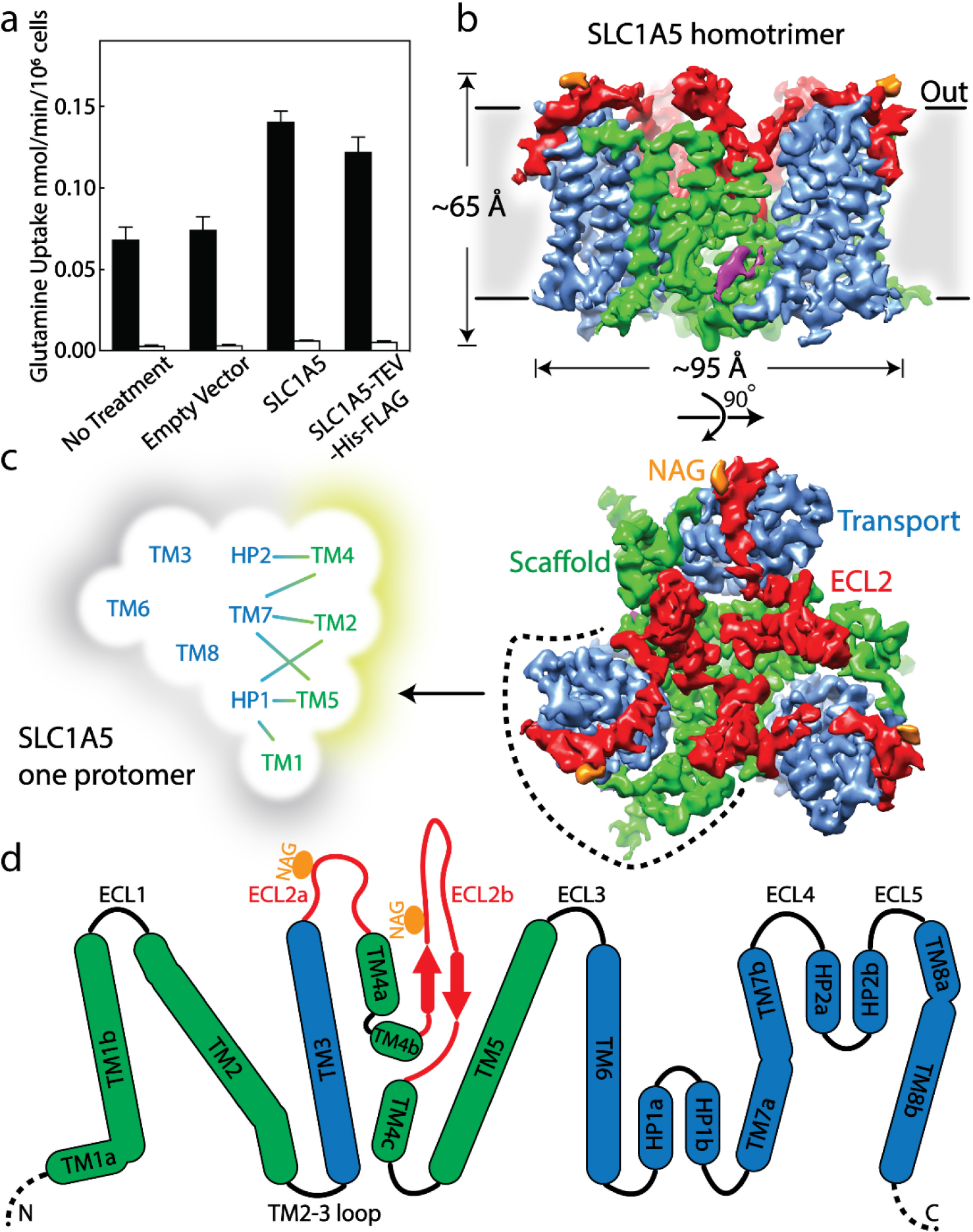
Cryo-EM structure of human SLC1A5 and its function. **a**, Uptake of [^14^C]-glutamine in HAP1 SLC1A5KO cells or SLC1A5KO cells transiently transfected with pcDNA3 vector containing the indicated sequence in the presence (solid bar) or absence (open bar) of sodium chloride. **b**, A density map of SLC1A5 homotrimer viewed from the side of the membrane (top) and the extracellular face (bottom) highlighting the scaffold domain (green), transport domain (navy), ECL2 (red) and N-acetyl-D-glucosamine (NAG, in orange). The density assigned to CHS is colored in magenta. **c,** Micelle/membrane and trimerization interfaces are highlighted in gray and wheat, respectively. Interactions between transport and scaffold domains are illustrated with lines. **d,** A schematic representation of the domains present in human SLC1A5. The N-linked glycosylation sites are indicated by orange circle. Dashed lines represent residues disordered in the structure.

The structure of SLC1A5 shows a homotrimer with an overall Glt_PH_-like fold in an outward-facing conformation, in contrast to the recently published inward-facing conformation of SLC1A5 (Fig. 1b) (7, 9, 15, 16). Comparison with all available outward-facing structures gave overall r.m.s.d. values of 1.2-2.3 Å, highlighting the similarity of these outward conformations (SI Appendix, Fig. S5, Table S2). Each SLC1A5 protomer contains two domains, a transport domain and a scaffold domain connected by the extracellular loop region 2 (ECL2a and ECL2b) (Fig. 1b-d, SI Appendix, Fig. S6). Superposition of all protomers reveals that three protomers from the outward-facing SLC1A5 trimer share a similar global fold. However, the deviation of the main chains upon the 3-fold operation indicates the outward-facing SLC1A5 trimers adopt the pseudo 3-fold symmetry (SI Appendix, Fig. S7). The individual transport domain containing transmembrane helices TM3, TM6-TM8 and helical loops 1-2 (HP1-HP2) interacts exclusively with the scaffold domain from its own protomer. The scaffold domain including TM1-TM2 and TM4-TM5 is involved in inter-protomer interactions and forms a compact core. The TM1, TM3 and TM6 make exclusive contacts with the detergent micelle and anchor the homotrimer in the membrane (Fig. 1c). The ECL2 connects TM3 to TM5 via TM4 and bridges the two domains. ECL2a spans a distance of about 55 Å across a quarter of the transport domain, sequentially interacting with ECL3, TM8, TM7a, and HP2b on extracellular surface. Based on sequence alignment, both SLC1A4 and SLC1A5 have the longest ECL2a region compared to other SLC1 family members including prokaryotic homologues (SI Appendix, Fig. S6).

Interestingly, the ECL2b exhibits extensive sequence divergence among SLC1 family members (SI Appendix, Fig. S6). ECL2b contains two short antiparallel β-strands with a long loop and protrudes into the extracellular surface (Fig. 1b, d) (16). Importantly, the ECL2b was reported to play a critical role in determining the receptor properties of SLC1A5 for retroviruses (20). Using an instrument configuration developed for the challenging intact mass analysis of membrane proteins (Methods), we were able to determine that SLC1A5 purified from GnTI-cells contains two oligo-mannose sites, consistent with the previously reported N-glycosylation sites Asn163 and Asn212 in ECL2 (SI Appendix, Fig. S8a) (21). Extra EM density close to Asn163 was clearly observed and modeled as an N-acetyl-D-glucosamine monomer (SI Appendix, Fig. S8b). The density corresponding to the N-glycosylation site on Asn212 was not well-defined.

Notably, a distinct cavity of ~550 Å^3^ was observed at the center of the homotrimer (SI Appendix, Fig. S9-10). The cavity is exclusively composed of hydrophobic residues and is filled with an unidentified, non-protein density in the cryo-EM map (SI Appendix, Fig. S9c). A similar cavity was also observed in Glt_PH_ and SLC1A3 structures (7, 15). Superposition of the inward-facing state of SLC1A5 (16) reveals a cavity in a similar location. The three helices of TM4 form significant inter-protomer contacts along the three-fold axis and are highly conserved in the SLC1 family, consistent with their critical structural role (SI Appendix, Fig. S10). The distinct hydrophobic interaction involving Phe201 is further stabilized by hydrogen-bonds between Arg202 and Glu227 from a neighboring protomer on the extracellular surface (SI Appendix, Fig. S10d).

### Outward- and Inward-facing transition of SLC1 family

Superposition with the inward-facing state demonstrates that the scaffold domain remains relatively rigid, while the transport domain moves toward the cytoplasm and the substrate binding site shifts by 19 Å toward the cytoplasmic face (Fig. 2a-b and SI Appendix, Fig. S5) (14–16, 19, 22). The ECL3 possessing one α-helix with ~1.5 turns connects the TM6 to the TM5 of the scaffold domain in the outward-facing state (Fig. 2a). In the inward-facing state (PDB: 6GCT), this 1.5 turn α-helix in ECL3 merges into TM6 which extends the length of TM6 by about 9.5 Å (Fig. 2a, c) (16). As a result, the shortened ECL3 further pulls the N-terminus of TM6 close to the TM5 causing the movement of TM6 toward the cytoplasm with ~40° tilt compared to the outward-facing state (Fig. 2a). Simultaneously, in order to compensate the movement of the transport domain and the conformational changes of TM6, the N-terminus of TM3 unwinds by approximately 1.5 turns and extends the TM2-3 loop in the cytoplasmic space (Fig. 2b-c). The extension of TM2-3 loop increases the distance between TM2 and TM3, resulting in a ~20° tilt compared to the outward-facing state (Fig. 2b). Furthermore, superimposition of all available structures shows similar structural changes between the outward- and inward-facing states (SI Appendix, Fig. S11a) (7–9, 11, 13, 14, 16, 18, 19, 22, 23). The structurally symmetrical TM3 and TM6 serve as a platform which holds the core region of the transport domain (Fig. 2c). During the transition between outward- and inward-facing states, the changes in lengths of the TM3, TM2-3 loop, TM6 and ECL3 regions trigger the adjustment in the platform plane relative to the membrane plane, resulting in the movement of the transport domain coupled with a ~30° rotation (Fig. 2a-c and 5) (16).

**Figure 2.**
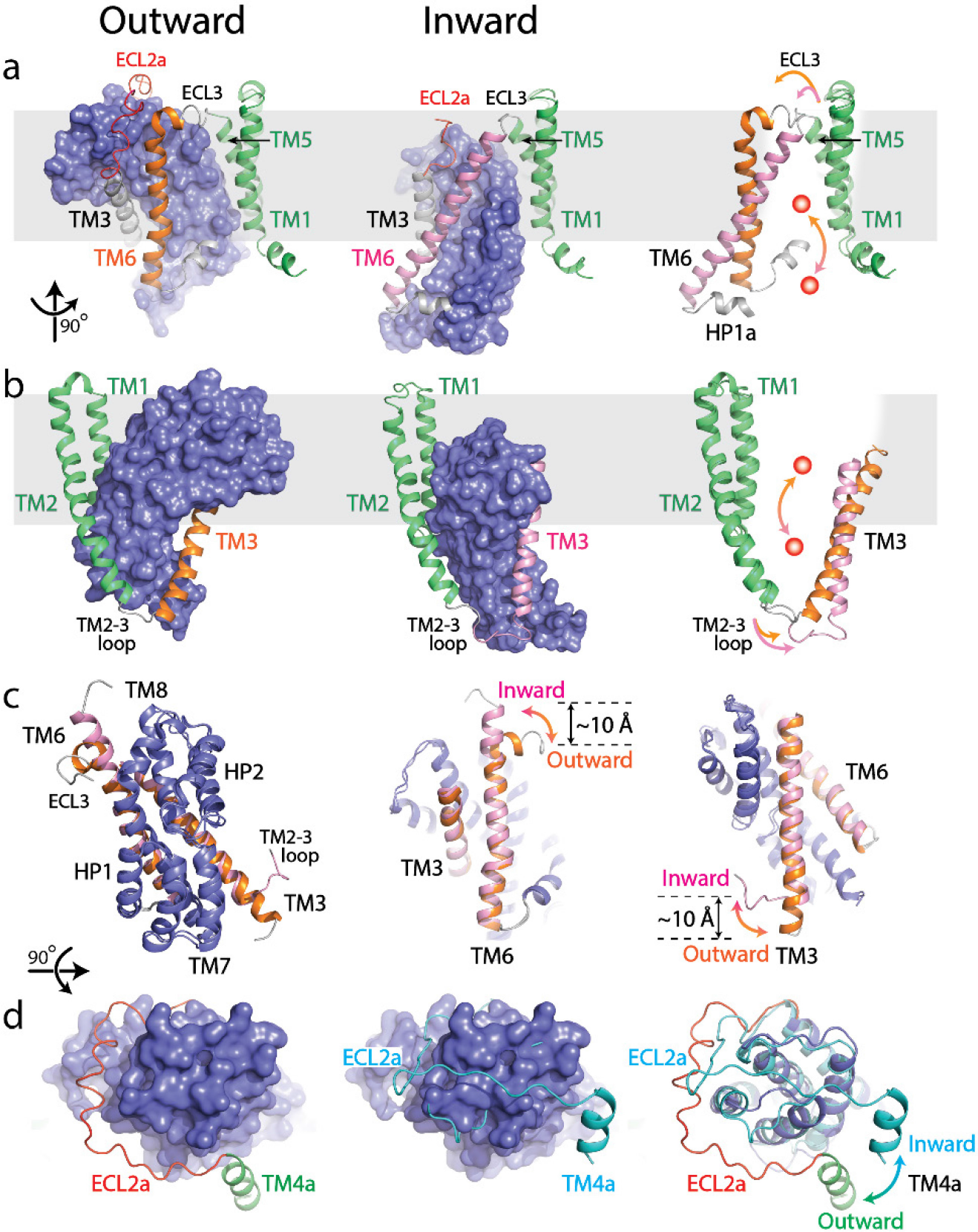
Structural comparison between outward- and inward-facing states of SLC1A5. **a and b**, Structure of the SLC1A5 monomer viewed from the side of the membrane highlighting TM6 and TM3, respectively. Transport domains are represented as molecular surfaces and colored in navy. Scaffold domains are in green. TM6 and TM3 are colored in orange in the outward-facing state (left) and pink in the inward-facing state (middle, PDB: 6GCT). Overlay of scaffold domains of inward- and outward-facing states highlighting TM3 and TM6 (right). The glutamine substrate is shown as a red ball. **c**, Superposition of the transport domains of SLC1A5 in the outward- and inward-facing (PDB: 6GCT) states (left). The conformational changes of TM6 (middle) and TM3 (right) between the outward-facing and inward-facing states are highlighted. **d**, Transport domains of SLC1A5 in an outward-facing state (left) and a cross-linked GltPH in inward-facing state (middle, PDB:3KBC) viewed from the extracellular face. The SLC1A5 transport domain in the outward-facing conformation serving as the reference is shown in molecular surface. The conformational changes of ECL2a between the outward- (in red) and inward-facing (in cyan) states are highlighted (right).

Notably, the well-defined ECL2a undergoes a large conformational rearrangement, swinging out by ~35° relative to the transport domain when compared with an inward-facing state of a cross-linked GltPH structure (Fig. 2d) (9). In the outward-facing state, the ECL2a encompasses one side of the transport domain and is positioned to expose both the ECL4 and ECL5. However, in the inward-facing state, the ECL2a forms a sharp turn and sits on top of the transport domain covering a narrow groove formed by the ECL4 and ECL5 to accommodate rigid body movement of the transport domain (9, 18, 19, 23). Overlaying all available structures shows distinct conformations on ECL2a (SI Appendix, Fig. S11b). In the outward-facing state, there are two observed conformational populations. One is represented by the ECL2a encircling the transport domain (PDB: 1XFH, 2NWX, 2NWW, 2NWL, 4IZM, 6BMI, 6BAU, 6BAVand 6BAT) as observed in our SLC1A5 cryo-EM structures (6MP6 and 6MPB) (7, 8, 22, 23). The other population is represented by the ECL2a burying ECL4 and ECL5 (PDB: 4OYE, 5CFY, 4OYF, 5E9S, 5DWY and 4KY0) (13, 14, 18). In the inward-facing state, all of ECL2a is repositioned to extend along the ECL4 and ECL5 (PDB: 3V8G, 3KBC, 3V8F, 4P6H, 4P19, 4P1A, 4P3J, 4X2S, 6GCT) (9, 11, 16, 18, 19). The pose of ECL2b does not change regardless of the conformational changes with an r.m.s.d of 1.5 Å (SI Appendix, Fig. S9a, S10b) (16).

The interface between the transport and scaffold domains buries a surface of ~ 2,300 Å^2^ in the outward-facing state. This interface is 1.7 times larger than that of the inward-facing state (Fig. 3a-c) (16). The domain interface in the two conformational states is dominated by hydrophobic amino acids. These hydrophobic residues in the scaffold domain emanate from almost the same set of residues, which are highly conserved among the SLC1 family (SI Appendix, Fig. S6). The scaffold domain forms major contacts with HP1 and the N-terminal region of TM7 (zone 2) in outward-facing state (Fig. 3a-b, SI Appendix, Fig. S6). In contrast, the scaffold domain interacts with HP2 and the N-terminal region of TM8 (zone 1) in the inward-facing state (Fig. 3c) (16). Interestingly, the transport domain exploits a different set of variable hydrophobic residues with similar hydrophobicity between the two states (Fig. 3a-c). Two conserved bulky hydrophobic residues (Phe377 and Trp461) of the transport domain are involved in both the outward- and inward-facing states, suggesting the crucial role of these residues in transport function (Fig. 3d, SI Appendix, Fig. S6). In the outward-facing state, ECL2a is likely to be situated at the side of the transport domain, partially occupying the zone 1 and it overlaps with TM2, and TM5 in the inward-facing state (Fig. 3e). In order to transition from the outward- to inward-facing state, the ECL2a has to be repositioned to fully expose zone 1.

**Figure 3.**
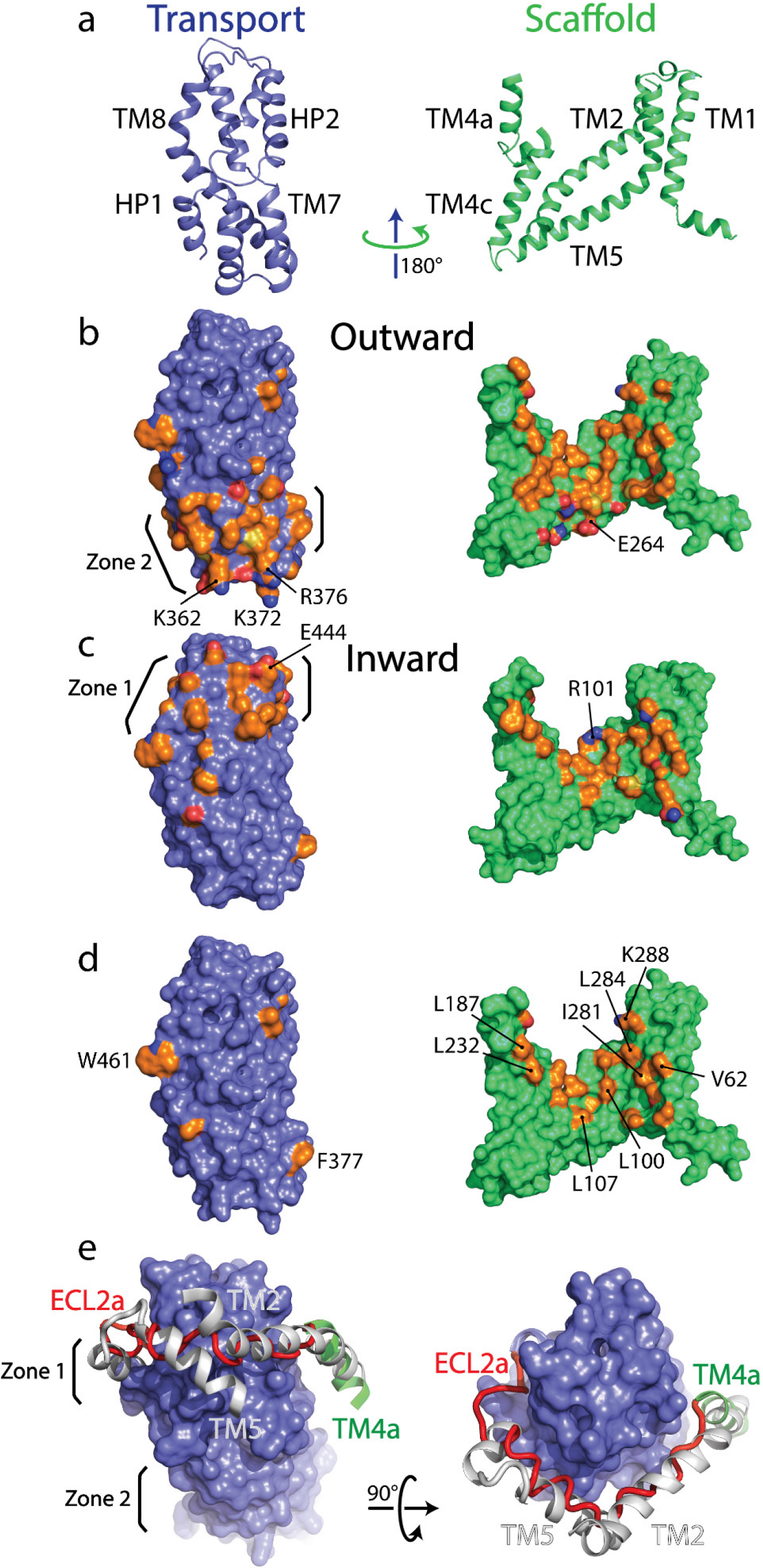
The interface between the scaffold and transport domain from SLC1A5 in the outward- and inward-facing states. **a.** One reference SLC1A5 model viewed from the membrane is shown in cartoon with the transport domain 180o rotated exposing the interface, with the secondary structural elements labeled. **b. c.** Molecular surface of the interface in the outward- and inward-facing state (PDB:6GCT). The interacting residues from the transport domain were clustered as Zone 2 and Zone 1, respectively. **d.** Molecular surface with the common interacting residues shared in both the inward- and outward-facing states. **e.** Superposition of the transport domains of inward- and outward-facing state showing key structural changes in ECL2. Using the transport domain as the reference, overlay of the ELC2a and TM4 (represented in red and green cartoon, respectively) shown in the outward-facing state with TM2, and TM5 (represented in gray cartoon) in the inward-facing state. The transport domain is shown in navy and scaffold domain is in green. The interface area is colored by the elements of the interface amino acids (carbon in orange, oxygen in red and nitrogen in blue). Common amino acids within the interface were labeled (see also SI Appendix, Fig. S6).

The extracellular and cytoplasmic surfaces of the transport domains show opposite electrostatic potential. Several acidic amino acid residues including Asp410, Glu444 and Asp451 are clustered on extracellular face of the transport domain. In contrast, basic residues including Lys362, Lys372 and Arg376 are clustered on the intracellular surface of the transport domain (Fig. 3b-c). The salt bridge between Glu444 of the transport domain and Arg101 of the scaffold domain was reported to be important for the stability of the inward-facing state (16). Similarly, Arg376 of the transport domain has a potential to form a salt bridge with the Glu264 of scaffold domain in the outward-facing structure (Fig. 3b-c). Both Arg376 and Glu264 are highly conserved among the SLC1 family (SI Appendix, Fig. S6).

### Glutamine binding site and substrate specificity

To understand the basis for substrate recognition by SLC1A5, we solved the cryo-EM structure of SLC1A5 in the presence of the substrate, L-Gln at 3.8 Å resolution (SI Appendix, Fig. S3). The density was sufficient to dock the L-Gln molecule (Fig. 4a). The Gln binding pocket is formed by two oppositely oriented reentrant loops (HP1 and HP2), TM7 and TM8 of the transport domain (Fig. 4a-b). The HP2 loop in the unliganded structure is in an open conformation providing a direct access to the binding site. Upon Gln binding, HP2 loop acts as a gatekeeper and comes within close proximity to the serine-rich HP1 loop to shield the substrate from the extracellular surface (Fig. 4a). Superposition of all available structures using transport domains revealed that the HP2 loop is closed upon substrate binding with HP1-2 tip distance of 4.5-6 Å regardless of conformational states (SI Appendix, Fig. S12, Table S2) (7–9, 11, 14–16, 18, 19, 22–24). In inhibitor-bound structures, the HP2 loops are largely either open or disordered (8, 15, 22). For structures in the absence of small molecule ligands, including sodium-bound and unbound states the HP1-2 tip distance varies (13, 14, 18). Furthermore, the substrate binding interaction is maintained between the two states of SLC1A5 (Fig. 4c) (16). Both α-amino and α-carboxylate groups of the substrate appear to be anchored by the carbonyl backbone of Ser351, the amide backbone and the sidechain of Ser353 in the HP1 loop and can be further stabilized by sidechains of Asp464 and Asn471 in TM8 (Fig. 4b). Importantly, the Nɛ2 of the substrate, L-Gln, is within hydrogen bond distance to the sidechain of Asp464 in TM8. A smaller substrate, L-Asn, could fit into the binding site to satisfy the interaction with Asp464, consistent with the comparable K_m_ values of human SLC1A5 for L-Asn and L-Gln determined in proteoliposomes (25). The position of L-Gln in our SLC1A5 structure is similar to that of the L-Asp in the SLC1A3 crystal structure, suggesting a conserved-ligand binding pocket among SLC1 family members (Fig. 4d) (7, 15).

**Figure 4.**
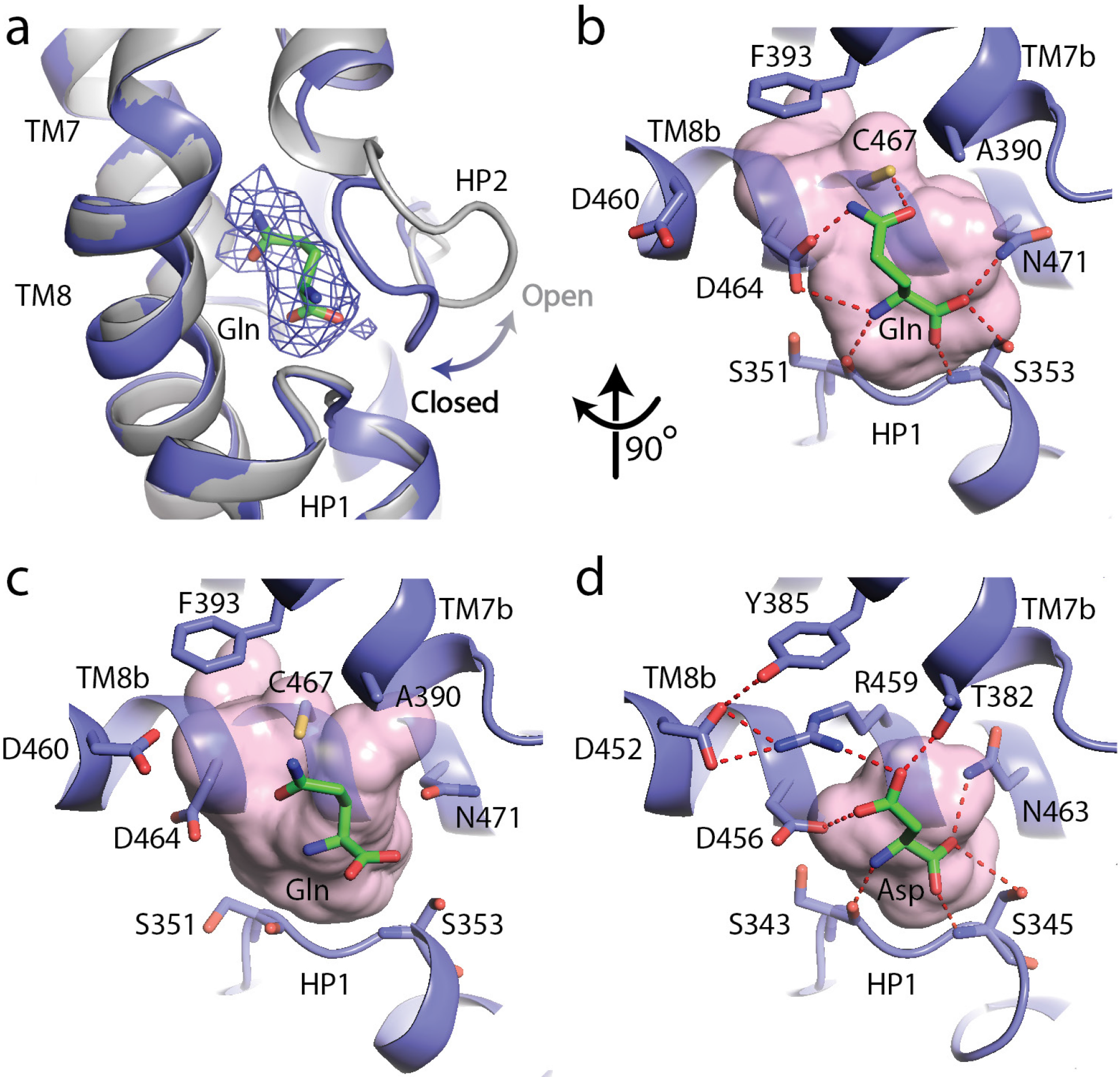
The substrate, L-glutamine-bound SLC1A5 structure. **a**, Conformational change for the HP2 upon binding of L-glutamine in the outward-facing state of SLC1A5. The unliganded structure is shown in grey and the L-glutamine-bound structure is in navy. EM density over the ligand is shown as a blue mesh. **b** and **c**, Zoomed in views of the substrate binding sites of the outward-facing and inward-facing states (PDB:6GCT) of SLC1A5, respectively. **d**, Interaction of L-Asp in the outward-facing state of SLC1A3 (PDB code: 5LLU). Addition of the substrate increases the ability of the transporter to transition between the conformations supported by increased dynamics in HDX (SI Appendix, Fig. S16). Critical side chains are labeled (stick representation) with hydrogen bonds (red dashed lines). Molecular surface in the substrate binding sites are in pink.

Cys467, a unique residue among the SLC1 family, is at a suitable distance to donate its proton to the carbonyl group of the Gln substrate (Fig. 4b, SI Appendix, Fig. S6) and could thus contribute to substrate selectivity. Mutation of Cys467 to Ala in SLC1A5 showed significant increase in K_m_ for Gln (26). The corresponding residue in all glutamate transporters of the SLC1 family is a conserved bulkier Arg residue (SI Appendix, Fig. S6). In SLC1A3, the Arg residue with its guanidinium group forms a hydrogen bond with the carboxylate group of the substrate Asp, and a favorable cation-*π* interaction with a Tyr residue (Fig. 4d). As a result of the smaller amino acid at this location, SLC1A5 has an additional side cavity involving Phe393, Val436, Ala433, Val463 and Asp464, whereas no such cavity is present in SLC1A3 (Fig. 4b, d). Besides the difference in the pocket size and shape, Arg in place of the Cys will lead to steric clashes with the Gln substrate in SLC1A5. The critical role of the Cys467 in SLC1A5 for substrate specificity was further supported by an equivalent mutation of threonine in SLC1A4 (Thr459) to a cysteine, resulting in alteration to L-Gln binding (22) and inhibition by Cys modifying reagents in proteoliposome assays (27). Ala390, a conserved residue in neutral amino acid transporters, SLC1A4 and SLC1A5, does not form any contact with the Gln substrate and could tolerate small neutral amino acid substrates (Fig. 4b, SI Appendix, Fig. S6). In contrast, the structurally equivalent residue Thr in SLC1A3 forms a hydrogen bond with the carboxylate group of the substrate.

### Cholesteryl hemisuccinate binding site

Stretches of residual EM densities were observed in a hydrophobic crevice near the domain interface. Based on the shape of the density and the presence of cholesteryl hemisuccinate (CHS) during the early steps of purification process, we assigned this density to CHS. This density was observed in cryo-EM structures of SLC1A5 in the presence or absence of substrate Gln (Fig. 1b). Using an unbiased lipidomics approach, CHS was detected as a major component in the sample used for cryo-EM. The identity of CHS was further confirmed by mass spectrometry using commercially available CHS as a reference (SI Appendix, Fig. S13). Although these data do not confirm CHS is directly bound to SLC1A5, the mass spectrometry data does confirm its presence in the sample prior to cryo-EM.

The cryo-EM density supports a specific binding site for CHS in the outward-facing SLC1A5 conformation. Electron density suggests that CHS binds inside a hydrophobic pocket formed by TM3, TM4a, TM4c and TM7 from one protomer, as well as TM5 from the adjacent protomer (SI Appendix, Fig. S14). The putative lipid density was also observed in the inward-facing state of SLC1A5, located in a hydrophobic pocket formed by TM3, TM4a, TM4c, and HP2a from one protomer, and TM5 from the adjacent protomer (16). Docking a CHS molecule in this putative density shows that the corresponding location is similar to that of the outward-facing state despite different pocket shape between two conformational states (SI Appendix, Fig. S14). Remarkably, the specific CHS-binding site in SLC1A5 overlaps with the UCPH_101_ allosteric binding site reported in SLC1A3 structure (15) (SI Appendix, Fig. S15). The highly lipophilic UCPH_101_ binds in the outward-facing conformation and blocks the movement of the transport domain relative to the scaffold domain. Additional putative lipid density was reported previously in the inward-facing state of SLC1A5, located between the transport and scaffold domains within the same protomer (16). The corresponding density was not observed in the outward-facing state of SLC1A5 due to differences in the local environment between the transport and scaffold domains upon the conformational changes.

### SLC1A5 solution dynamics

Hydrogen/deuterium exchange (HDX) on SLC1A5 reveals a very rigid scaffold domain with low (less than 10%) deuterium exchange in TM helices, similar to that of TM helices for GPCRs (28) (SI Appendix, Fig. S16a). In contrast, the transport domain shows moderate deuterium exchange (10-50%) indicating this is the more dynamic domain. The addition of L-Gln imparted conformational changes to five regions of the transporter (SI Appendix, Fig. S16b). TM4a/loop (188-196), ECL4 (401-404), HP2 (421-439) and ECL5 (446-455) displayed increased conformational mobility (increased exchange) in the presence of L-Gln, whereas TM7a (381-384) showed decreased conformational mobility (decreased exchange). Addition of the substrate L-Gln could increase the frequency of switching between outward- and inward-facing conformations, which likely accounts for the increased conformational mobility observed in the HDX experiment. This would also suggest that the closed conformation of HP2 in our structure represents one of the conformations involved in shuttling L-Gln through the transporter. The decreased conformational mobility at TM7a (381-384) could be indicative of a direct or allosteric interaction with L-Gln. Alternatively, this could suggest the location for a sodium binding site if L-Gln enhances sodium affinity for SLC1A5. Interestingly, a sodium ion was resolved in this location in the SLC1A3 structure (15). Although the resolution of the structure here did not enable detection of sodium, the HDX protection at this site would support a model for enhanced sodium affinity in the presence of L-Gln.

## Discussion

Human SLC1A5 is the first eukaryotic sodium-dependent neutral amino acid transporter for which both the inward- and outward-facing conformational states have been mapped by cryo-EM. Superposition using the transport domains from the Gln-bound outward-facing and inward-facing (16) cryo-EM structures shows similar poses at the substrate binding site at this resolution (Fig. 4b, c). However, in the outward-facing state, the transport domain presents a larger interface to the scaffold domain compared to the inward-facing state. In order to compensate for the loss of the binding energy, a much larger non-polar surface (~1,000 Å^2^ for SCL1A5 trimer) is imbedded into the lipid membrane in the inward-facing state. Burial of the non-polar surface into the lipid membrane may help facilitate conformational plasticity for the transporter. Further structural studies of unbound SLC1A5 in the inward-facing state will be needed to fully understand the molecular drivers for the structural transition.

Our cryo-EM structures of unbound and substrate-bound SLC1A5 revealed that HP2 is the extracellular gate that controls entry of the substrate (8, 15). As a gatekeeper, HP2 in a closed conformation comes within close proximity to the serine-rich HP1, shielding the substrate from the extracellular surface upon substrate binding (Fig. 4a). The structure of the unliganded form of SLC1A5 provides an example with the HP2 loop in an open conformation and serves as a useful starting model for designing competitive inhibitors such as TBOA (8, 29). The opening of the HP2 loop increases the accessible surface area, exposing a previously unidentified cryptic pocket which could accommodate a large inhibitor that would be specific to the HP2 open state. In contrast, the closed conformation of HP2 has a small binding site, limiting the size of the inhibitors (30, 31).

The human SLC1A5 represents a distinct SLC1 subfamily identifiable in part by the Cys and Ala residues for substrate specificity and the unique conformation of the two ECL2a and ECL2b loops allowing for structural flexibility at TM4b for the key trimer interaction. Furthermore, ECL2a encircles the transport domain in the outward-facing state, covering a large part of the interaction surface for the inward facing state. In the inward-facing state the ECL2a is repositioned, exposing the interaction surface for the scaffold domain (Fig. 5) (16).

**Figure 5.**
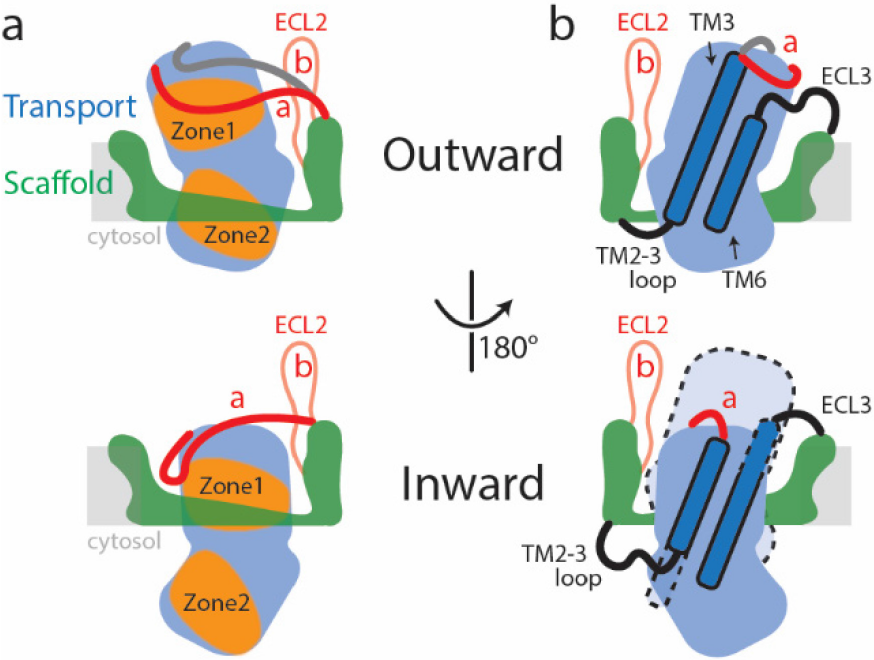
Schematic representation of conformational changes in SLC1A5 during substrate transport. **a**. In the outward-facing state (top), the Zone 2 from the transport domain interacts with the scaffold domain, with the Zone 1 partially occupied by the ECL2a. Two different poses of the ECL2a were observed (shown in red and dark grey lines): one is situated at the side of the transport domain (red line); the other is crossing over the crests of ECL4 and ECL5 (dark grey line) (SI Appendix, Fig. S11b). In the inward-facing state (bottom), the ECL2a has to be repositioned, releasing the Zone 1 to interact with the scaffold domain. **b**. The inherent plasticity around the TM2-3 loop, ECL3, TM3, and TM6 regions (highlighted in dash lines) allows the transitions between the outward-facing (top) and inward-facing states (bottom). The relative position of transport domain in the outward-facing state is highlighted in dash (bottom panel in b). The regions of SLC1A5 are colored based on the Fig. 1b. The light grey line depicts membrane boundary.

The existence of a potential allosteric binding pocket seen in the outward- and inward-facing (16) SLC1A5 structures could facilitate the design of selective inhibitors that can bind to a site that is remote from the conserved substrate-binding site. Such allosteric druggable pockets at the lipid-exposed surface near the intracellular portions of membrane proteins have recently been structurally elucidated in several proteins and this raises new opportunities for drug design (SI Appendix, Fig. S15). We believe that these findings provide new and exciting structural insights and opportunities for antagonizing glutamine metabolism at the transporter level. Our structures of human SLC1A5, in both unbound and substrate-bound forms, represent the first high resolution cryo-EM structures of SLC1 family captured in the outward-facing state. These structures are an early example of the value that cryo-EM can bring to structure-based drug design.

## Materials and Methods

The gene encoding SLC1A5 was cloned into pcDNA3.1^+^ (Life Technologies) where it was fused to a C-terminal His10-Flag affinity double tag separated by TEV protease cleavage site. The protein was expressed in HEK293S GnTI^−^ cells (ATCC CRL-3022) grown in Expi293 Expression Media (Life Technologies) to densities 2.5 × 10^6^ cells ml^−1^. Cells were transiently transfected in Opti-MEM reduced serum media (Life Technologies) using ExpiFectamine Transfection Kit (Life Technologies). Cells were collected at 48 h after transfection. MAb cKM4012 was cloned and expressed, as previously reported with a small modification (32, 33). Detailed procedures for assays, cryo-EM image collection and structural analysis are described in SI Appendix, Materials and Methods.

## Acknowledgements

We thank James Smith, Chia-Chin Lee, Elizabeth Dushin and Kelsey Oleynek for assisting with cloning, expression and protein purification of SLC1A5. We also thank Mark Tibbitts for cell transfection. Y.Y.D. and E.P.C. are members of the SGC, (Charity ref: 1097737) funded by AbbVie, Bayer Pharma AG, Boehringer Ingelheim, the Canada Foundation for Innovation, Genome Canada, GlaxoSmithKline, Janssen, Lilly Canada, Merck & Co., Novartis, the Ontario Ministry of Economic Development and Innovation, Pfizer, São Paulo Research Foundation-FAPESP and Takeda, as well as the Innovative Medicines Initiative Joint Undertaking ULTRA-DD grant 115766 and the Wellcome Trust106169/Z/14/Z. We thank Kingsley Sumner for IT support and Ravi Kurumbail for critical reading.

