## Supplemental Table 1 for "Structural basis for the transport mechanism of the human glutamine transporter SLC1A5 (ASCT2)"

Name: Seungil Han

Address: Pfizer Inc., Eastern Point Road, Groton, CT 06340

### 1. Materials and Methods

**SLC1A5 and SLC1A5_cKM4012 (Fab) complex purification.** Cells were resuspended at 1:10 (w/v) ratio in 25 mM HEPES pH 7.0, 300 mM NaCl, 0.2 mM Tris (2-carboxyethyl) phosphine (TCEP), buffer supplemented with EDTA free protease inhibitor cocktail (Roche) and 2.5 µl/ml of Benzonase nuclease (Sigma), and disrupted in Microfluidizer processor M110L (Microfluidics) at 12,000 psi. Cell lysate was clarified by centrifugation at 4,000*g* for 30 min, and membrane fraction was collected by ultracentrifugation at 225,000*g* for 60 min. Membrane pellets were homogenized at 1:5 (w/v) ratio in 25 mM HEPES pH 7.0, 300 mM NaCl, 1% lauryl maltose neopentyl glycol (LMNG) (Anatrace), 0.1% cholesterol hemisuccinate (CHS) (Anatrace), 0.2 mM TCEP buffer followed by 2h solubilization with gentle agitation at 4 °C. Insoluble material was removed by ultracentrifugation at 290,000*g* for 60 min. Solubilized material was incubated overnight at 4 °C with anti-flag M2 affinity agarose resin (Sigma) with gentle agitation. 1 ml of the resin efficiently captured solubilized SLC1A5 from 10g of cells. Unbound material was removed by centrifugation at 1,000*g* for 10 min and the resin was washed with 25 mM HEPES pH 7.0, 300 mM NaCl, 0.003% LMNG, 0.2 mM TCEP buffer (buffer A) three times. SLC1A5 was eluted with buffer A supplemented with 0.2 mg/ml Flag peptide (CPC Scientific) and concentrated to 1 mg $ml^{-1}$ using Amicon Ultra Ultracell-100K (Millipore Sigma) centrifugal filter. The affinity purified protein was applied to Superose 6 Increase 10/300 gel filtration column (GE Healthcare) equilibrated with buffer A. All protein purification steps performed on ice or at 4 °C at all times.

Fab fragments of cKM4012 monoclonal antibodies were generated using the Fab Preparation Kit (Pierce) and further purified on Superdex 75 column equilibrated with 30 mM HEPES pH 7.0, 100 mM NaCl, 0.003% LMNG, 10 mM L-glutamine (L-Gln). SLC1A5/cKM4012 (Fab)/L-Gln complex was formed in the followed order: 0.85 ml of SLC1A5 at 16 µM was mixed with 0.05 ml of L-Gln at 200 mM to achieve 10 mM final L-Gln concentration, incubated 20 min in ice, mixed with 1 ml of cKM4012 Fab fragments at 16 µM, incubated 20 min in ice, concentrated to 0.1 ml on Amicon Ultra Ultracell-100K (Millipore Sigma) centrifugal filter. The complex was loaded to Superose 6 10/300 Increase filtration column (GE Healthcare) equilibrated with 25 mM HEPES pH 7.0, 100 mM NaCl, 0.003% LMNG, 10 mM L-Gln. Two top peak fractions were taken and concentrated to 2 mg/ml.

**L-Glutamine Uptake Assay.** HAP1 SLC1A5 knock out (SLC1A5KO) cells (Horizon Discovery HZGHC005452c002) contain a 1bp insertion in a coding exon of SLC1A5 and do not express SLC1A5 (as described by the vendor and confirmed by us). SLC1A5KO cells were transiently transfected by static electroporation using a MaxCyte STX system. Viable cells ($10^6$ per mL) were transfected with 1.0 μg of pcDNA3.1 (empty vector) or with 1 μg of either pcDNA3.1-FL-SLC1A5 or pcDNA3.1-FL-SLC1A5-TEV-His-Flag using the OC-100 or OC-400 processing assembly (electroporation cell) and conditions specified by the vendor. Transfected cells were then cultured at 37 °C in a humidified environment in 5% carbon dioxide. After 24 hrs, transfected cells were removed flasks and frozen for later use. Glutamine uptake was determined by measuring the incorporation of radio-labeled glutamine into SLC1A5KO cells transfected with the indicated DNA. Specifically, frozen SLC1A5KO cells were thawed then cultured overnight in T-175 flasks in Iscove's modified Dulbecco's medium (Gibco 12440) supplemented with 10 % fetal bovine serum. At the time of assay, the cells were removed from flasks with cell dissociation buffer (Gibco 13151), centrifuged at 800 x g, re-suspended in assay buffer (25 mM Tris pH 7.4, 2 mM KCl, 1 mM MgCl2, 1 mM CaCl2, 5 mM glucose with or without and 100 mM NaCl) and plated at 50,000 cells per well in a 384-well Cytostar-T scintillation microplate (PerkinElmer RPNQ0166). The plate was then incubated at 37 °C in a humidified environment in 5% carbon dioxide. After ~ 60 minutes, the plate was removed from the incubator and placed at room temperature (RT). After ~15 minutes, the assay was initiated by the addition of a mixture of unlabeled glutamine and $^{14}$C-labeled glutamine (PerkinElmer NEC4150) prepared in assay buffer. The plate was then covered with a plastic seal and [$^{14}$C]-glutamine uptake was determined at 40 minutes by scintillation counting with a MicroBeta2 (2450 Microplate Counter; Perkin Elmer). The final assay volume was 40 μL and the final assay conditions were 25 mM TRIS pH 7.4, 2 mM KCl, 1 mM $MgCl_2$, 1 mM $CaCl_2$, 5 mM glucose, with or without 100 mM NaCl and 30 μM glutamine/[$^{14}$C]-glutamine (50 dpm/pmol).

**Grid preparation and data acquisition.** 3.5 μL of 2.0 mg/ml purified SLC1A5_cKM4012 (Fab) with/without L_Gln complex was applied to the glow-discharged Quantifoil Au R1.2/1.3 grid (Structure Probe), and subsequently vitrified using a Vitrobot Mark IV (FEI Company). In order to overcome an orientation bias, n-octyl-β -d-glucopyranoside (BOG, Anatrace) was added to the sample prior freezing. Cryo grids were loaded into a Titan Krios transmission electron microscope (ThermoFisher Scientific) operating at 300 keV with a Gatan K2 Summit direct electron detector. Images were recorded with SerialEM in super-resolution mode with a super resolution pixel size of 0.543 Å and a defocus range of 1.2 to 2.5 μm. Data were collected with a dose rate of 5 electrons per physical pixel per second, and images were recorded with a 10s exposure and 250 ms subframes (40 total frames) corresponding to a total dose of 42 electrons per $Å^2$. All details corresponding to individual datasets are summarized in Table S1.

**Electron microscopy data processing.** For SLC1A5-cKM4012 (Fab) complex, a total of 3,804 dose-fractioned movies were gain-corrected, 2 x binned (resulting in a pixel size of 1.086 Å), and beam-induced motion correction using MotionCor2 (1) with the dose-weighting option. The SLC1A5_cKM4012 (Fab) particles were automatically picked from the dose-weighted, motion corrected average images using Gautomatch. CTF parameters were determined by Gctf (2). A total of 743,525 particles were then extracted using Relion 2.0 (3) with a box size of 224 pixels. The 2D, 3D classification and refinement were performed with Relion 2.0. Two rounds of 2D classification and one round of 3D classification were performed to select the homogenous particles. 3D classification results showed only the outward-facing state with the substrate binding site accessible from the extracellular side. After selecting particle coordinates, per-particle CTF estimation was refined using the program Gctf (2). One set of 170,751 particles was then submitted to 3D auto-refinement. All 3D classifications and 3D refinements were started from a 60 Å low-pass filtered version of an ab initio map generated with VIPER (4). The particles were re-centered using the refined particle offsets before being re-extracted, followed by parameter conversion and final Auto_Refine in cisTEM. Auto_Refine in cisTEM, using a soft binary mask that removed the micelle and Fab regions, yielded the final reconstruction at 3.5 Å global resolution, improving the resolution by ~0.3 Å. The global resolution was evaluated using conventional Fourier Shell Correlation analysis. Prior to visualization, all density maps were sharpened by applying different negative temperature factors (-15, -30, and -50 $Å^2$) using automated procedures (5), along with the half maps, were used for model building. Local resolution was determined using ResMap (6) (Figure S2). Data collection, processing and refinement of SLC1A5-cKM4012 (Fab) with L-Gln substrate complex were performed similarly as described for SLC1A5-cKM4012 (Fab) (Figure S3). Fab signals subtraction followed by refinements with/without C3 symmetry operation did not improve the resolutions for both the SLC1A5-cKM4012 (Fab) and SLC1A5-cKM4012 (Fab)_Gln complexes. The reason could be the Fab-bound SLC1A5 is perhaps pseudo-3 fold symmetric. The presence of the Fab disrupts the low-resolution pseudo-symmetry and facilitates particle alignment. However, the flexibility of the Fabs harms the high-resolution reconstruction at the SLC1A5 trimer regions. A soft binary mask was applied to the SLC1A5 trimer region during the last several rounds of Auto_Refine in cisTEM to improve the local resolution.

**Model building and refinement** The initial template of the human SLC1A5 was derived from a homology-based model calculated by SWISS-MODEL (7). The model was docked into the EM density map using Chimera (8) and followed by manually adjustment using COOT (9). For the SLC1A5_L-Gln modeling, one L-glutamine molecular was generated and manually docked into the density using COOT. Each model was independently subjected to global refinement and minimization in real space using the module phenix.real_space_refine in PHENIX(10) against separate EM half-maps with default parameters. The model was refined into a working half-map, and improvement of the model was monitored using the free half map. The geometry parameters of the final models were validated in Coot and using MolProbity and EMRinger (11). These refinements were performed iteratively until no further improvements were observed. The final refinement statistics were provided in Table S1. Model overfitting was evaluated through its refinement against one cryo-EM half map. FSC curves were calculated between the resulting model and the working half map as well as between the resulting model and the free half and full maps for cross-validation (Figure S4). Figures were produced using PyMOL (12) and Chimera (8).

**SLC1A5 proteoliposomes production.** Liposomes were prepared according to Pingitore *et al.* with modifications (13). A 9:1 mixture of L-α-phosphatidylcholine (PC) (Sigma) and 18:1 biotinyl cap phosphoethanolamine (PE) (Avanti) was dried under $N_2$ gas stream, rehydrated in water by vigorous mixing on a vortex mixer followed by extrusion through 400 nm polycarbonate membranes (Sigma) resulting in 6.7% liposomes. SLC1A5 proteoliposomes were formed during detergent-mediated reconstitution. First, pre-made liposomes were mixed with C12E8 to final 1.4% of liposomes and 1.7% of C12E8 in 20 mM Tris, pH 7.0, 10 mM L-Gln followed by 30 minutes incubation at RT. Second, purified SLC1A5 in 20 mM Tris, pH 7.0, 100 mM 100 NaCl, 10 mM L-Gln, 0.05% n-dodecyl-β-D-maltopyranoside (DDM) and 6 mM β-mercaptoethanol was added to lipid-detergent-micellar solution to make 2000:1 lipid to protein ratio. The protein-lipid-detergent mixture was incubated 30 min at RT before detergents were removed through absorption on hydrophobic resin Amberlite XAD-4. To exchange to an uptake buffer, proteoliposomes were passed through the 7K MWCO Zeba spin desalting columns (ThermoFisher Scientific) equilibrated with 20 mM Tris, pH 7.0, 15 mM Sucrose to balance internal osmolality.

**L-glutamine uptake assay using proteoliposomes**. L-glutamine uptake was determined by measuring the incorporation of radio-labeled L-glutamine into biotinylated-SLC1A5 proteoliposomes. Specifically, the assay was initiated by the addition of biotinylated-SLC1A5 proteoliposomes into triplicate wells of a 96-well polypropylene plate with each well containing the indicated concentration of NaCl and 50 $\mu$M L-glutamine/ L-[3,4-$^3$H)]-glutamine (100 dpm/pmol). After the indicated time at room temperature (~22 ºC), the assay was stopped by the addition of 15 µl of ice-cold streptavidin beads (Pierce™ Streptavidin Agarose) prepared in 20 mM Tris, pH 7.0, 15 mM sucrose buffer. After 10 min at room temperature the biotin proteoliposome/streptavidin bead complex was transferred to a 96-well GF/C filter plate (PerkinElmer) plate (pre-blocked with 0.1% BSA and washed x 2 in 20 mM Tris, pH 7.0, 15 mM sucrose buffer). The plate was then vacuum filtered and then washed three times with 500 µl per well of ice-cold 20 mM Tris, pH 7.0, 15 mM sucrose buffer. The plate was then air dried overnight and 50 *u*l of Betaplate Scint (PerkinElmer) was added to each well of the 96-well plate. The plate was then sealed and glutamine uptake was determined by scintillation counting with a MicroBeta2 (PerkinElmer).

**Intact protein mass spectrometry.** Purified SLC1A5 was diluted to 0.1 mg/mL in MilliQ water just prior to analysis. 5 µl was injected into an Agilent 1290 UPLC system and was separated using a Agilent PLRP-S column (1000Å pore size, 5.0 *u*m particle size, 50 x 2.1 mm). The protein was separated using a linear gradient from 0 to 100% acetonitrile in 0.1% formic acid. The sample analysis was carried out on an Agilent 6530 QTof mass spectrometer equipped with a Dual AJS electrospray source operated in positive ion mode. Raw mass spectra were viewed using MassHunter (version B.07.00 Service Pack 2, Agilent) and mass spectral deconvolution was performed using BioConfirm (B.07.00, Agilent).

**Cholesteryl hemisuccinate (CHS) detection by untargeted LC/MS.** Purified SLC1A5 protein sample was treated with 10X acetonitrile for 1 h to denature proteins. Precipitated proteins were pelleted using centrifugation at 15000 x g for 10 min and the supernatant was transferred to a new vial. The supernatant was dried completely using a speed-vac and resuspended with 100 µl of 2:2:1 isopropanol:acetonitrile:water. CHS was detected using high resolution mass spectrometer (Q-Exactive plus) in series with an Agilent 1290 UPLC pump. We modified the waters application note: Lipid Separation using UPLC with Charged Surface Hybrid Technology. Briefly, the mobile phase system consisted of (A) acetonitrile: water (60:40) with 10 mM ammonium formate and 0.1% formic acid and (B) isopropanol:acetonitrile (90:10) with 10 mM ammonium formate and 0.1% formic acid. Metabolites were separated using an Acquity UPLC CSH C18 2.1 x 100 mm, 1.7 µm column heat to 55 ºC. 10 µL of each sample were subjected to the following gradient: time 0 = 40% B, time 2 = 43% B, time 2.1 = 50% B, time 12 = 54% B, time 12.1 = 70% B, time 18 = 99% B, time 18.1 = 40% B, time 20 = 40% B. The mass spectrometer was set with the following parameters: mass range = 100 – 1500 m/z, negative mode, 70,000 resolution, AGC target = 1e6, Maximum IT = 100 ms, Centroid, ddMS2 resolution = 30,000, AGC target 1e5, Maximum IT 50 ms, Top N =5, Loop count = 5, Isolation window = 4.0 m/z, CE = 30. The data was processed using Compound Discoverer 2.0.

**Hydrogen/Deuterium exchange mass spectrometry (HDX-MS).** HDX experiments on SLC1A5 were carried out at 4 °C using a similar system to those previously described (14). HDX studies used the same SLC1A5 construct used for CryoEM, except the final stock storage buffer was at pH 8.5. Compounds were added to a 12uM solution of SLC1A5 at a concentration of 200uM. In brief, 5 ul of the 12µM SLC1A5 solution (with or without ligand) was incubated in a $D_2O$ containing buffer (50 mM HEPBS, pH 8.5, 150 mM NaCl, 2% (v/v) glycerol, 0.003% (m/v) LMNG) for six exchange times (10 s, 30 s, 1 m, 5 m, 15 m, and 1 h) before quenching the deuterium exchange reaction with an acidic quench solution (pH 2.4). All mixing and digestions were carried out on a LEAP Technologies PAL liquid handling robot housed inside a chromatographer's refrigerator. Digestion was performed in-line with chromatography using an immobilized pepsin column (NovaBioAssay). Mass spectra were acquired on a Velos Pro mass spectrometer (ThermoFisher Scientific) and peptide identification from the MSMS data was done using Mascot. Deuterium exchange values for peptide isotopic envelopes at each time point were calculated and processed using the Workbench software (15).

### 2. Additional Data

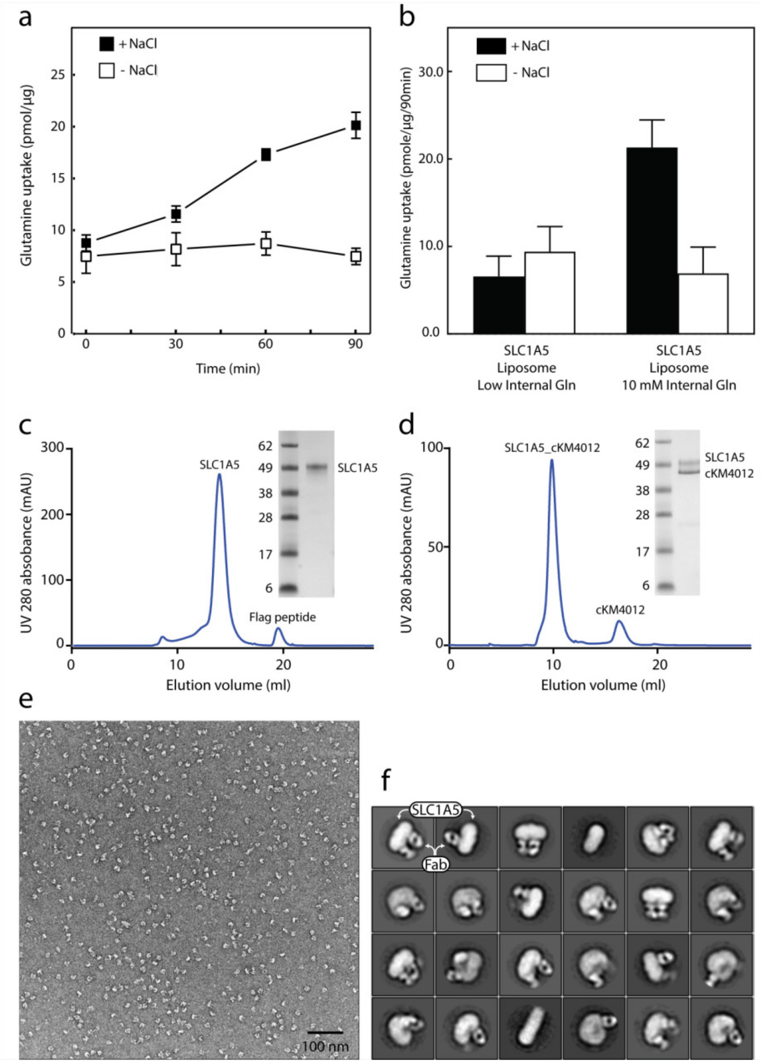

**Figure S1 | SLC1A5 protein purification, functional assay in proteoliposomes and negative staining. a**, **b**, Uptake of [$^3$H]-L-glutamine by purified SLC1A5 reconstituted in proteoliposomes. Transport was started by the addition of 50 µM L-glutamine/[3H]-L-glutamine (100 dpm/pmole) to SLC1A5 proteoliposomes in the presence (▪) or absence (▫) of 100 mM external sodium chloride. **a**, Internal proteoliposome buffer contained 10 mM glutamine and uptake was measured at the indicated times. The experiment was performed with triplicates at each time point; error bars represent standard deviation. **b**, Internal buffer contained either low internal glutamine (0 mM or 0.1 mM) or 10 mM glutamine and uptake was measured at 90 min. The graphs are means of three independent experiments performed in triplicate. Error bars represent the s.e.m. Similarly prepared SLC1A5 in proteoliposomes showed [$^3$H]-glutamine efflux in the presence of external (assay buffer) glutamate and NaCl (16). **c**, **d**, Size-exclusion chromatography (Superose 6 10/300 Increase filtration column) of SLC1A5 and its complex with cKM4012 (Fab). The peak fraction was examined by SDS-polyacrylamide gel electrophoresis (SDS-PAGE). **e**, Raw micrograph of SLC1A5-cKM4012 (Fab) complex by negative-stain electron microscopy. **f**, 2D-class averages of particles from negative-stain electron microscopy of SLC1A5-cKM4012 (Fab) complex. Signals corresponding to the SLC1A5 and Fabs are highlighted.

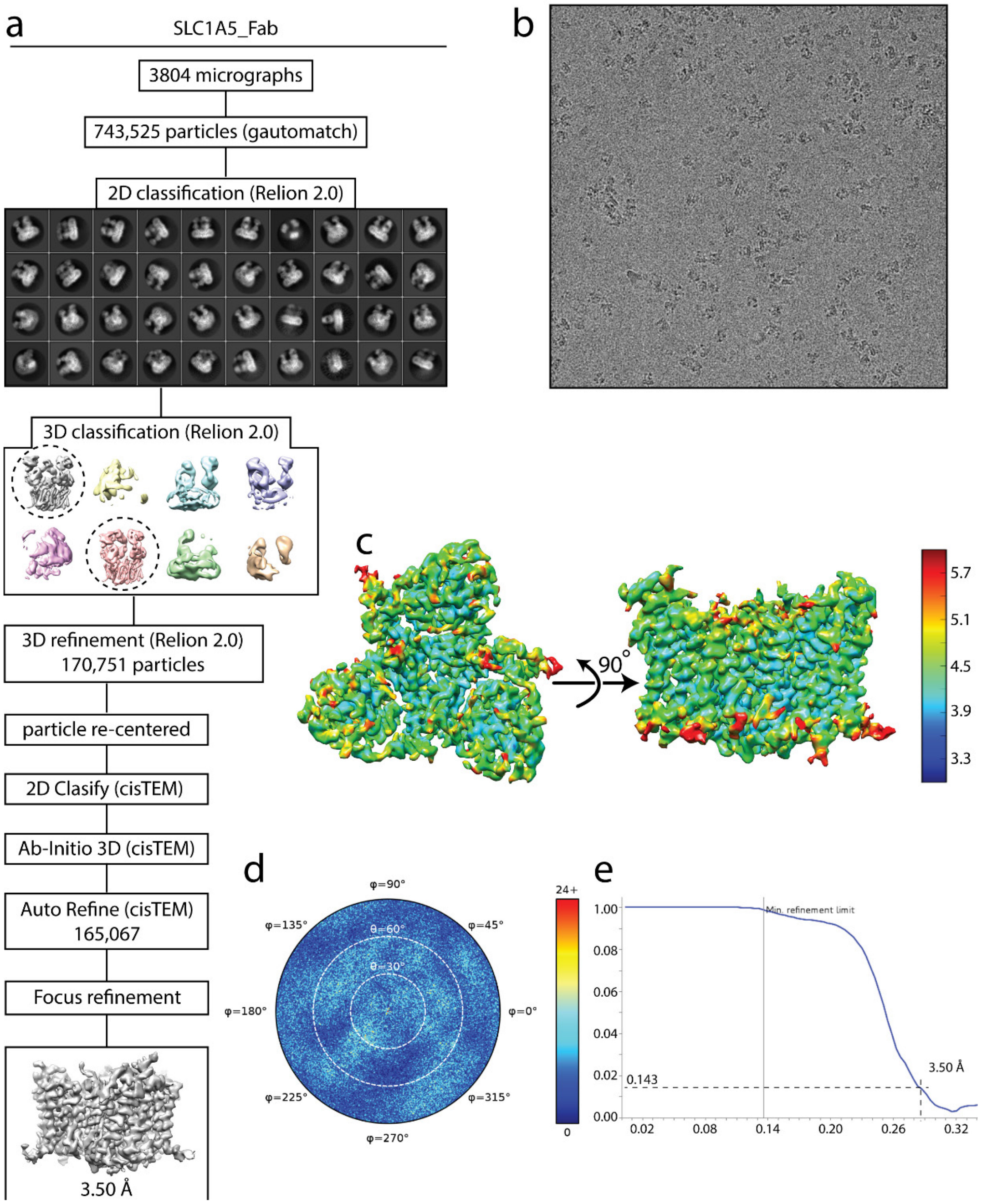

**Figure S2 | Cryo-EM analysis of SLC1A5-cKM4012 (Fab) complex a**, Flow chart of the cryo-EM data processing procedure. Details can be found in the Methods. **b**, A representative cryo-EM micrograph. **c**, Local resolution of the map estimated using the ResMap program and colored as indicated. **d,** Angular orientation distribution of the particles used in the final reconstruction. The particle distribution is indicated by different color shades. **e**, Fourier shell correlation (FSC) curve of the structure with FSC as a function of resolution using cisTEM output. The resolution is ~3.5 Å at the FSC cutoff of 0.143. A thin vertical line indicates that only spatial frequencies to 1/ (7.8 Å) are used to determine particle alignment parameters during refinement in cisTEM.

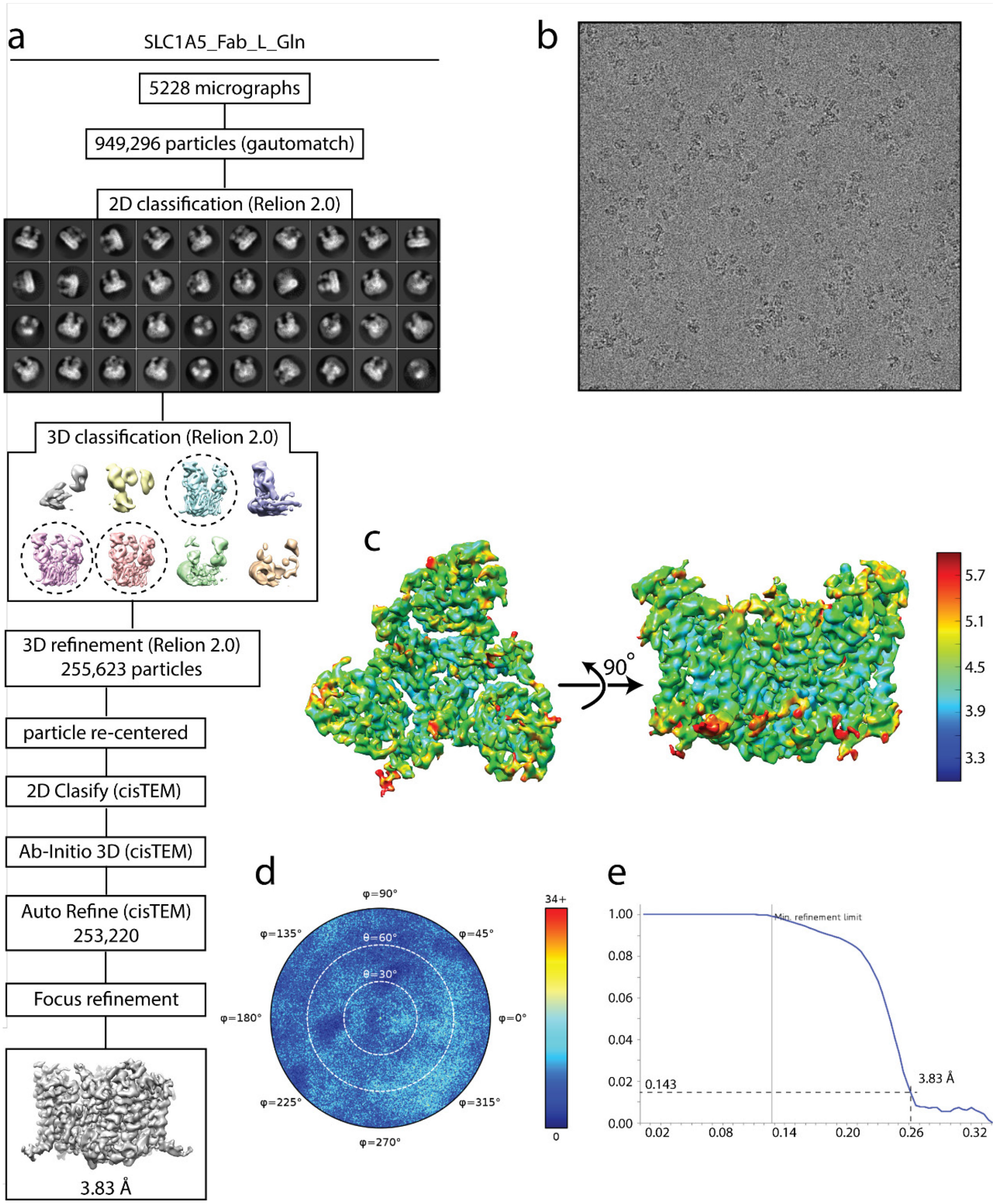

**Figure S3 | Cryo-EM analysis of SLC1A5-cKM4012 (Fab) complex in the presence of L-glutamine. a**, Flow chart of the cryo-EM data processing procedure. Details can be found in the Methods. **b**, A representative cryo-EM micrograph. **c**, Local resolution of the map estimated using the ResMap program and colored as indicated. **d**, Angular orientation distribution of the particles used in the final reconstruction. The particle distribution is indicated by different color shades. **e**, Fourier shell correlation (FSC) curve of the structure with FSC as a function of resolution using cisTEM output. The resolution is ~3.8 Å at the FSC cutoff of 0.143. A thin vertical line indicates that only spatial frequencies to 1/ (7.8 Å) are used to determine particle alignment parameters during refinement in cisTEM.

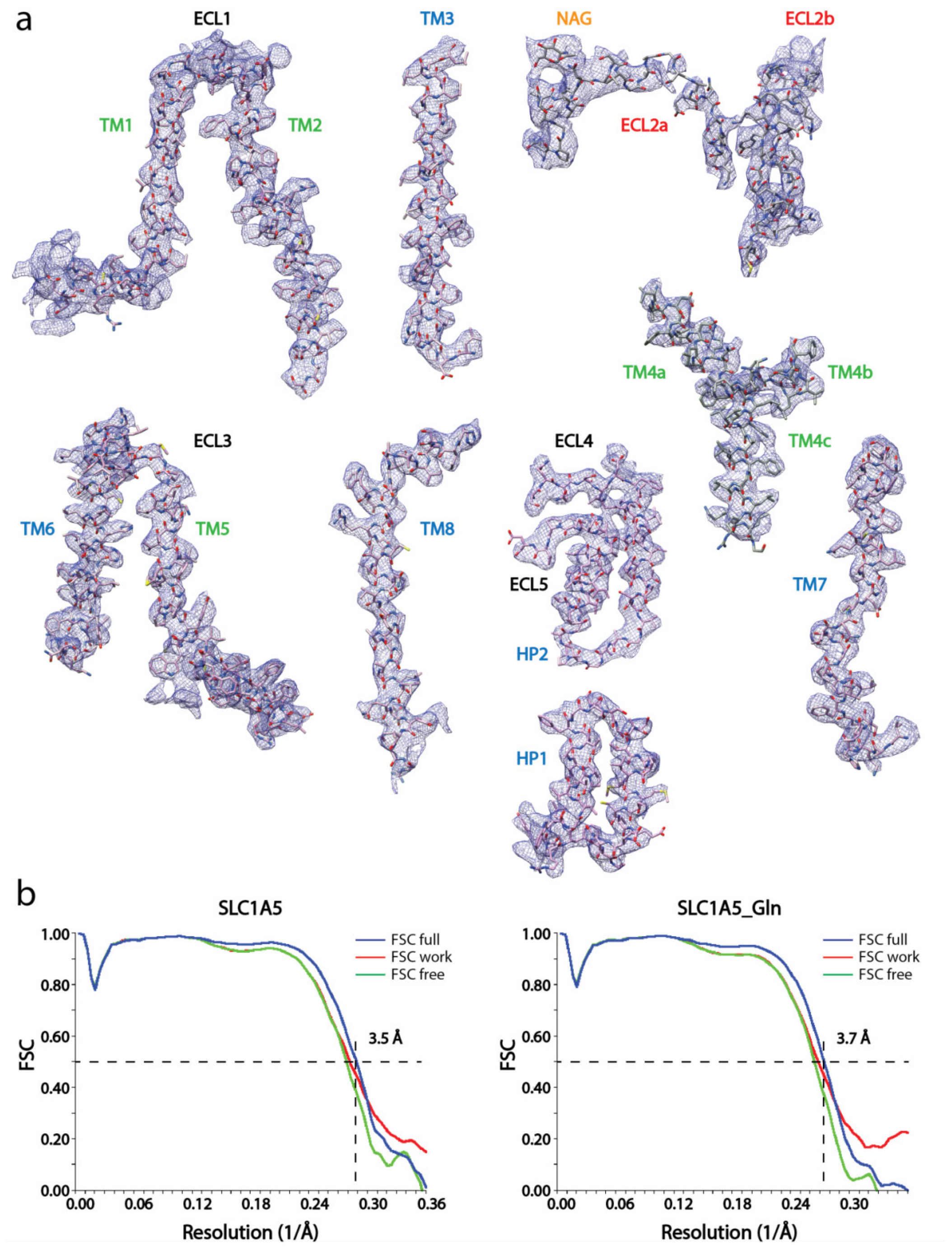

**Figure S4 | Cryo-EM densities of the eight transmembrane helices with ECL loops of SLC1A5-cKM4012 (Fab). a**, Cryo-EM density is sharpened with a negative b-factor 15 Å$^2$ and displayed at the contour level 8σ for the transmembrane helixes and loops regions, except for the ECL2a, and b regions at 5.6σ and 4σ respectively. The atomic model with side chains are shown as sticks. The segments are labeled and colored as in Figure 1c. **b**, Model validation. Comparison of the FSC curves between model and half map 1 (work), model and half map 2 (free), and model and full map are plotted in red, green and blue, respectively.

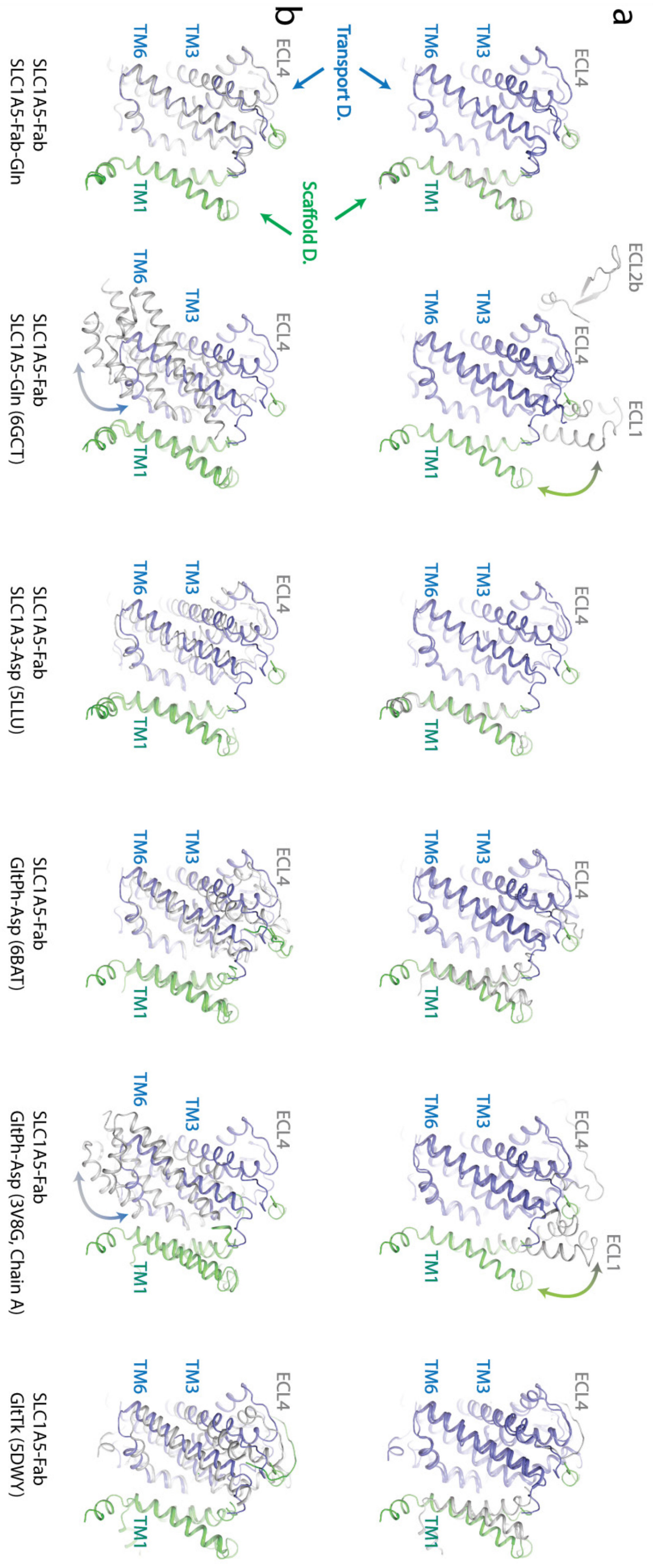

a
b
Transport D.
Scaffold D.
ECL4
ECL2b
ECL1
TM3
TM6
TM1
SLC1A5-Fab
SLC1A5-Fab-Gln
SLC1A5-Fab
SLC1A5-Gln (6GCT)
SLC1A5-Fab
SLC1A3-Asp (5LLU)
SLC1A5-Fab
GltPh-Asp (6BAT)
SLC1A5-Fab
GltPh-Asp (3V8G, Chain A)
SLC1A5-Fab
GltTk (5DWY)

**Figure S5 | Superposition of the SLC1A5-cKM4012 (Fab) structure with other representive structures.** From left to right: SLC1A5-cKM4012 (Fab) with/without Gln structures; SLC1A5-cKM4012 (Fab) and SLC1A5 inward-facing structures; SLC1A5-cKM4012 (Fab) and SLC1A3 outward facing structures; SLC1A5-cKM4012 (Fab) and $Glt_{PH}$ outward-facing structures; SLC1A5-cKM4012 (Fab) and $Glt_{PH}$ inward-facing structures; SLC1A5-cKM4012 (Fab) and $Glt_{TK}$ outward-facing structures. All the structures are represented in cartoon. The transport and scaffold domains of SLC1A5-cKM4012 (Fab) are colored in slate and green, respectively. **a**, Superposition of the transport domain as a reference. The transport and scaffold domains of the representative structures are colored in slate and gray. **b**, Superposition of the scaffold domain as a reference. The transport and scaffold domains from the represented structures are colored in gray and green, respectively.

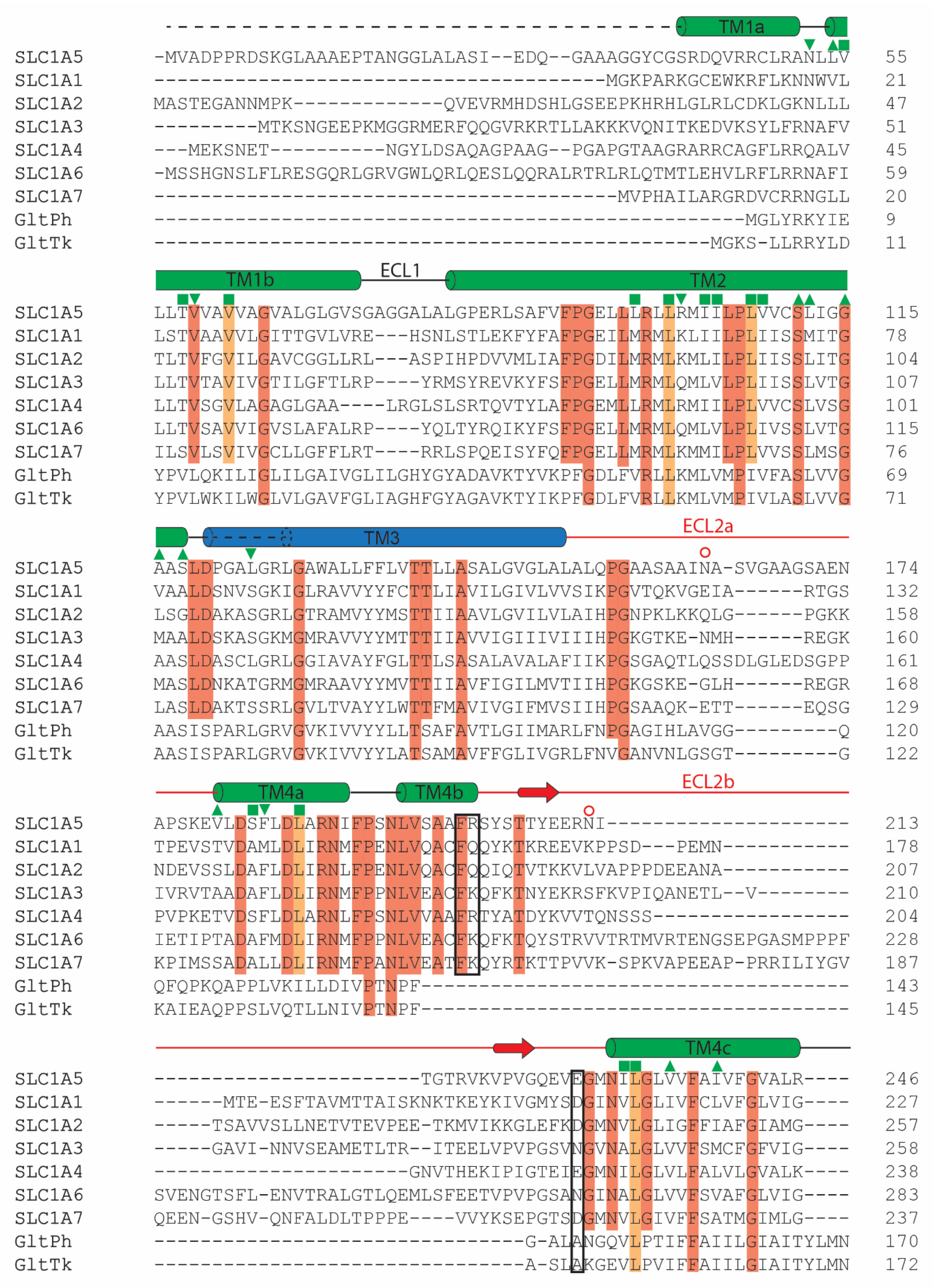
TM1a
SLC1A5 -MVADPPRDSKGLAAAEPTANGGLALASI--EDQ--GAAAGGYCGSRDQVRRCLRANLLV 55
SLC1A1 ---------------------------------------MGKPARKGCEWKRFLKNNWVL 21
SLC1A2 MASTEGANNMPK-------------QVEVRMHDSHLGSEEPKHRHLGLRLCDKLGKNLLL 47
SLC1A3 ---------MTKSNGEEPKMGGRMERFQQGVRKRTLLAKKKVQNITKEDVKSYLFRNAFV 51
SLC1A4 ---MEKSNET----------NGYLDSAQAGPAAG--PGAPGTAAGRARRCAGFLRRQALV 45
SLC1A6 -MSSHGNSLFLRESGQRLGRVGWLQRLQESLQQRALRTRLRLQTMTLEHVLRFLRRNAFI 59
SLC1A7 ----------------------------------------MVPHAILARGRDVCRRNGLL 20
GltPh ---------------------------------------------------MGLYRKYIE 9
GltTk ------------------------------------------------MGKS-LLRRYLD 11
TM1b
ECL1
TM2
SLC1A5 LLTVVAVVAGVALGLGVSGAGGALALGPERLSAFVFPGELLLRLLRMIILPLVVCSLIGG 115
SLC1A1 LSTVAAVVLGITTGVLVRE---HSNLSTLEKFYFAFPGEILMRMLKLIILPLIISSMITG 78
SLC1A2 TLTVFGVILGAVCGGLLRL---ASPIHPDVVMLIAFPGDILMRMLKMLILPLIISSLITG 104
SLC1A3 LLTVTAVIVGTILGFTLRP----YRMSYREVKYFSFPGELLMRMLQMLVLPLIISSLVTG 107
SLC1A4 LLTVSGVLAGAGLGAA----LRGLSLSRTQVTYLAFPGEMLLRMLRMIILPLVVCSLVSG 101
SLC1A6 LLTVSAVVIGVSLAFALRP----YQLTYRQIKYFSFPGELLMRMLQMLVLPLIVSSLVTG 115
SLC1A7 ILSVLSVIVGCLLGFFLRT----RRLSPQEISYFQFPGELLMRMLKMMILPLVVSSLMSG 76
GltPh YPVLQKILIGLILGAIVGLILGHYGYADAVKTYVKPFGDLFVRLLKMLVMPIVFASLVVG 69
GltTk YPVLWKILWGLVLGAVFGLIAGHFGYAGAVKTYIKPFGDLFVRLLKMLVMPIVLASLVVG 71
TM3
ECL2a
SLC1A5 AASLDPGALGRLGAWALLFFLVTTLLASALGVGLALALQPGAASAAINA-SVGAAGSAEN 174
SLC1A1 VAALDSNVSGKIGLRAVVYYFCTTLIAVILGIVLVVSIKPGVTQKVGEIA------RTGS 132
SLC1A2 LSGLDAKASGRLGTRAMVYYMSTTIIAAVLGVILVLAIHPGNPKLKKQLG------PGKK 158
SLC1A3 MAALDSKASGKMGMRAVVYYMTTTIIAVVIGIIIVIIIHPGKGTKE-NMH------REGK 160
SLC1A4 AASLDASCLGRLGGIAVAYFGLTTLSASALAVALAFIIKPGSGAQTLQSSDLGLEDSGPP 161
SLC1A6 MASLDNKATGRMGMRAAVYYMVTTIIAVFIGILMVTIIHPGKGSKE-GLH------REGR 168
SLC1A7 LASLDAKTSSRLGVLTVAYYLWTTFMAVIVGIFMVSIIHPGSAAQK-ETT------EQSG 129
GltPh AASISPARLGRVGVKIVVYYLLTSAFAVTLGIIMARLFNPGAGIHLAVGG---------Q 120
GltTk AASISPARLGRVGVKIVVYYLATSAMAVFFGLIVGRLFNVGANVNLGSGT---------G 122
TM4a
TM4b
ECL2b
SLC1A5 APSKEVLDSFLDLARNIFPSNLVSAAFRSYSTTYEERNI--------------------- 213
SLC1A1 TPEVSTVDAMLDLIRNMFPENLVQACFQQYKTKREEVKPPSD---PEMN----------- 178
SLC1A2 NDEVSSLDAFLDLIRNLFPENLVQACFQQIQTVTKKVLVAPPPDEEANA----------- 207
SLC1A3 IVRVTAADAFLDLIRNMFPPNLVEACFKQFKTNYEKRSFKVPIQANETL--V-------- 210
SLC1A4 PVPKETVDSFLDLARNLFPSNLVVAAFRTYATDYKVVTQNSSS----------------- 204
SLC1A6 IETIPTADAFMDLIRNMFPPNLVEACFKQFKTQYSTRVVTRTMVRTENGSEPGASMPPPF 228
SLC1A7 KPIMSSADALLDLIRNMFPANLVEATFKQYRTKTTPVVK-SPKVAPEEAP-PRRILIYGV 187
GltPh QFQPKQAPPLVKILLDIVPTNPF------------------------------------- 143
GltTk KAIEAQPPSLVQTLLNIVPTNPF------------------------------------- 145
TM4c
SLC1A5 -----------------------TGTRVKVPVGQEVEGMNILGLVVFAIVFGVALR---- 246
SLC1A1 ------MTE-ESFTAVMTTAISKNKTKEYKIVGMYSDGINVLGLIVFCLVFGLVIG---- 227
SLC1A2 -----TSAVVSLLNETVTEVPEE-TKMVIKKGLEFKDGMNVLGLIGFFIAFGIAMG---- 257
SLC1A3 -----GAVI-NNVSEAMETLTR--ITEELVPVPGSVNGVNALGLVVFSMCFGFVIG---- 258
SLC1A4 ----------------------GNVTHEKIPIGTEIEGMNILGLVLFALVLGVALK---- 238
SLC1A6 SVENGTSFL-ENVTRALGTLQEMLSFEETVPVPGSANGINALGLVVFSVAFGLVIG---- 283
SLC1A7 QEEN-GSHV-QNFALDLTPPPE----VVYKSEPGTSDGMNVLGIVFFSATMGIMLG---- 237
GltPh --------------------------------G-ALANGQVLPTIFFAIILGIAITYLMN 170
GltTk --------------------------------A-SLAKGEVLPVIFFAIILGIAITYLMN 172

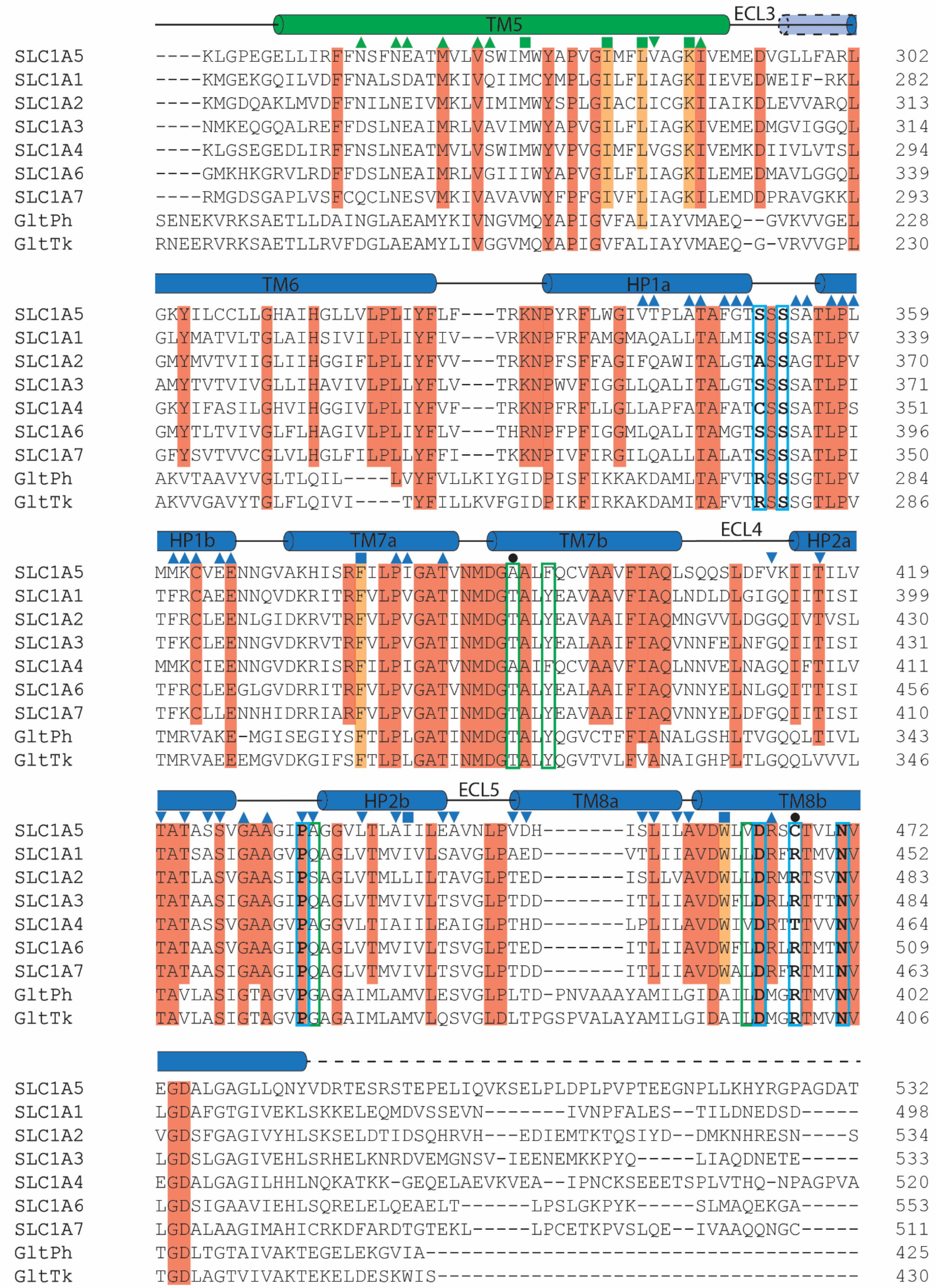
TM5
ECL3
SLC1A5 ----KLGPEGELLIRFFNSFNEATMVLVSWIMWYAPVGIMFLVAGKIVEMEDVGLLFARL 302
SLC1A1 ----KMGEKGQILVDFFNALSDATMKIVQIIMCYMPLGILFLIAGKIIEVEDWEIF-RKL 282
SLC1A2 ----KMGDQAKLMVDFFNILNEIVMKLVIMIMWYSPLGIACLICGKIIAIKDLEVVARQL 313
SLC1A3 ----NMKEQGQALREFFDSLNEAIMRLVAVIMWYAPVGILFLIAGKIVEMEDMGVIGGQL 314
SLC1A4 ----KLGSEGEDLIRFFNSLNEATMVLVSWIMWYVPVGIMFLVGSKIVEMKDIIVLVTSL 294
SLC1A6 ----GMKHKGRVLRDFFDSLNEAIMRLVGIIIWYAPVGILFLIAGKILEMEDMAVLGGQL 339
SLC1A7 ----RMGDSGAPLVSFCQCLNESVMKIVAVAVWYFPFGIVFLIAGKILEMDDPRAVGKKL 293
GltPh SENEKVRKSAETLLDAINGLAEAMYKIVNGVMQYAPIGVFALIAYVMAEQ--GVKVVGEL 228
GltTk RNEERVRKSAETLLRVFDGLAEAMYLIVGGVMQYAPIGVFALIAYVMAEQ-G-VRVVGPL 230
TM6
HP1a
SLC1A5 GKYILCCLLGHAIHGLLVLPLIYFLF---TRKNPYRFLWGIVTPLATAFGTSSSSATLPL 359
SLC1A1 GLYMATVLTGLAIHSIVILPLIYFIV---VRKNPFRFAMGMAQALLTALMISSSSATLPV 339
SLC1A2 GMYMVTVIIGLIIHGGIFLPLIYFVV---TRKNPFSFFAGIFQAWITALGTASSAGTLPV 370
SLC1A3 AMYTVTVIVGLLIHAVIVLPLLYFLV---TRKNPWVFIGGLLQALITALGTSSSSATLPI 371
SLC1A4 GKYIFASILGHVIHGGIVLPLIYFVF---TRKNPFRFLLGLLAPFATAFATCSSSATLPS 351
SLC1A6 GMYTLTVIVGLFLHAGIVLPLIYFLV---THRNPFPFIGGMLQALITAMGTSSSSATLPI 396
SLC1A7 GFYSVTVVCGLVLHGLFILPLLYFFI---TKKNPIVFIRGILQALLIALATSSSSATLPI 350
GltPh AKVTAAVYVGLTLQIL----LVYFVLLKIYGIDPISFIKKAKDAMLTAFVTRSSSGTLPV 284
GltTk AKVVGAVYTGLFLQIVI----TYFILLKVFGIDPIKFIRKAKDAMITAFVTRSSSGTLPV 286
HP1b
TM7a
TM7b
ECL4
HP2a
SLC1A5 MMKCVEENNGVAKHISRFILPIGATVNMDGAALFQCVAAVFIAQLSQQSLDFVKIITILV 419
SLC1A1 TFRCAEENNQVDKRITRFVLPVGATINMDGTALYEAVAAVFIAQLNDLDLGIGQIITISI 399
SLC1A2 TFRCLEENLGIDKRVTRFVLPVGATINMDGTALYEAVAAIFIAQMNGVVLDGGQIVTVSL 430
SLC1A3 TFKCLEENNGVDKRVTRFVLPVGATINMDGTALYEALAAIFIAQVNNFELNFGQIITISI 431
SLC1A4 MMKCIEENNGVDKRISRFILPIGATVNMDGAAIFQCVAAVFIAQLNNVELNAGQIFTILV 411
SLC1A6 TFRCLEEGLGVDRRITRFVLPVGATVNMDGTALYEALAAIFIAQVNNYELNLGQITTISI 456
SLC1A7 TFKCLLENNHIDRRIARFVLPVGATINMDGTALYEAVAAIFIAQVNNYELDFGQIITISI 410
GltPh TMRVAKE-MGISEGIYSFTLPLGATINMDGTALYQGVCTFFIANALGSHLTVGQQLTIVL 343
GltTk TMRVAEEEMGVDKGIFSFTLPLGATINMDGTALYQGVTVLFVANAIGHPLTLGQQLVVVL 346
HP2b
ECL5
TM8a
TM8b
SLC1A5 TATASSVGAAGIPAGGVLTLAIILEAVNLPVDH-------ISLILAVDWLVDRSCTVLNV 472
SLC1A1 TATSASIGAAGVPQAGLVTMVIVLSAVGLPAED-------VTLIIAVDWLLDRFRTMVNV 452
SLC1A2 TATLASVGAASIPSAGLVTMLLILTAVGLPTED-------ISLLVAVDWLLDRMRTSVNV 483
SLC1A3 TATAASIGAAGIPQAGLVTMVIVLTSVGLPTDD-------ITLIIAVDWFLDRLRTTTNV 484
SLC1A4 TATASSVGAAGVPAGGVLTIAIILEAIGLPTHD-------LPLILAVDWIVDRTTTVVNV 464
SLC1A6 TATAASVGAAGIPQAGLVTMVIVLTSVGLPTED-------ITLIIAVDWFLDRLRTMTNV 509
SLC1A7 TATAASIGAAGIPQAGLVTMVIVLTSVGLPTDD-------ITLIIAVDWALDRFRTMINV 463
GltPh TAVLASIGTAGVPGAGAIMLAMVLESVGLPLTD-PNVAAAYAMILGIDAILDMGRTMVNV 402
GltTk TAVLASIGTAGVPGAGAIMLAMVLQSVGLDLTPGSPVALAYAMILGIDAILDMGRTMVNV 406
SLC1A5 EGDALGAGLLQNYVDRTESRSTEPELIQVKSELPLDPLPVPTEEGNPLLKHYRGPAGDAT 532
SLC1A1 LGDAFGTGIVEKLSKKELEQMDVSSEVN-------IVNPFALES--TILDNEDSD----- 498
SLC1A2 VGDSFGAGIVYHLSKSELDTIDSQHRVH---EDIEMTKTQSIYD--DMKNHRESN----S 534
SLC1A3 LGDSLGAGIVEHLSRHELKNRDVEMGNSV-IEENEMKKPYQ-----LIAQDNETE----- 533
SLC1A4 EGDALGAGILHHLNQKATKK-GEQELAEVKVEA--IPNCKSEEETSPLVTHQ-NPAGPVA 520
SLC1A6 LGDSIGAAVIEHLSQRELELQEAELT------LPSLGKPYK-----SLMAQEKGA----- 553
SLC1A7 LGDALAAGIMAHICRKDFARDTGTEKL-----LPCETKPVSLQE--IVAAQQNGC----- 511
GltPh TGDLTGTAIVAKTEGELEKGVIA------------------------------------- 425
GltTk TGDLAGTVIVAKTEKELDESKWIS------------------------------------ 430

```
                  ---------
SLC1A5     --VASEKESVM-------------------------------------     541
SLC1A1     KKSYVNGGFAVD----KSDTISFTQTSQF-------------------     524
SLC1A2     QCVYAAHNSVIVDECKVTLA---------ANGKSADCSVEEEPWKREK     574
SLC1A3     KPI-DS-ETKM-------------------------------------     542
SLC1A4     APELESKESVL-------------------------------------     532
SLC1A6     RGRGGN-ESAM-------------------------------------     564
SLC1A7     KSVAEASELTLGPTCPHHVPVQVEQDEELPAASLNHCTIQISELETNV     560
GltPh      ------------------------------------------------     425
GltTk      ------------------------------------------------     430
```

**Figure S6 | Sequence alignment of human SLC1 transporters and two prokaryotic homologues.** The secondary structural features are presented according to the human SLC1A5 cryo-EM structures in this study. The color-coding scheme is the same as in Figure 1**d**. The secondary element changes on TM3 and TM6 upon outward- and inward-transitions are indicated. Conserved residues are highlighted and filled in red. Residues involved in the substrate binding and the neighboring pocket for SLC1A5 are boxed in cyan and green, respectively. Key residues for substrate specificity are indicated by black filled circles. The residues involved in inter-protomer interactions are boxed in black. Residues from Zone 2 (Figure 3) and their interacting residues from the scaffold domain in the inward-facing state are highlighted using ▼ and ▼, respectively. Residues from Zone 1 (Figure 3) and their interacting residues from the scaffold domain in the outward-facing state are highlighted using ▲, and ▲, respectively. Shared residues from the transport and scaffold domains during conformational changes are highlighted using ■ and ■, respectively. The conserved residues are filled in orange. The N-linked glycosylation sites are indicated by orange open circles. Dashed lines represent the sequences which are not observed from cryo-EM density.

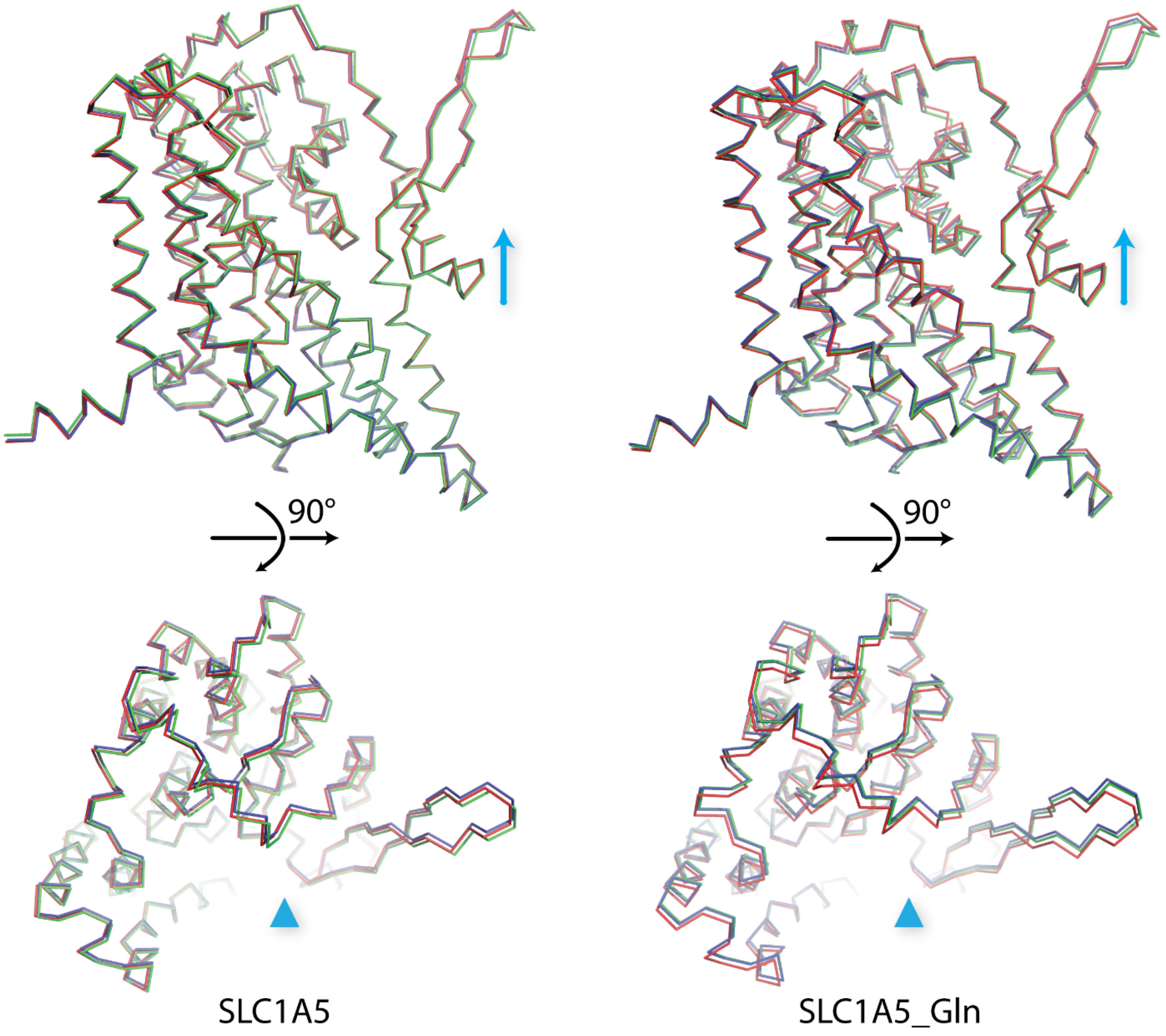

**Figure S7 | Pseudo 3-fold symmetry of outward-facing SLC1A5 structure.** SLC1A5 structures (unbound and Gln-bound) were repositioned using the scaffold domains of inward-facing SLC1A5 structure as the reference (PDB: 6GCT). Chain A and C were rotated around the z-axis (in cyan) by 120° and -120°, respectively, overlaying with the chain B. Three protomers from the outward-facing SLC1A5 strimmer were shown in ribbon and colored in green, blue, and red, respectively. Two views from the membrane (up) and cytoplasm (down) were shown. The results revealed that three protomers from the outward-facing SLC1A5 trimer share a similar global fold. However, the deviation of the main chains upon the 3-fold operation indicates the outward-facing SLC1A5 trimers adopt the pseudo 3-fold symmetry.

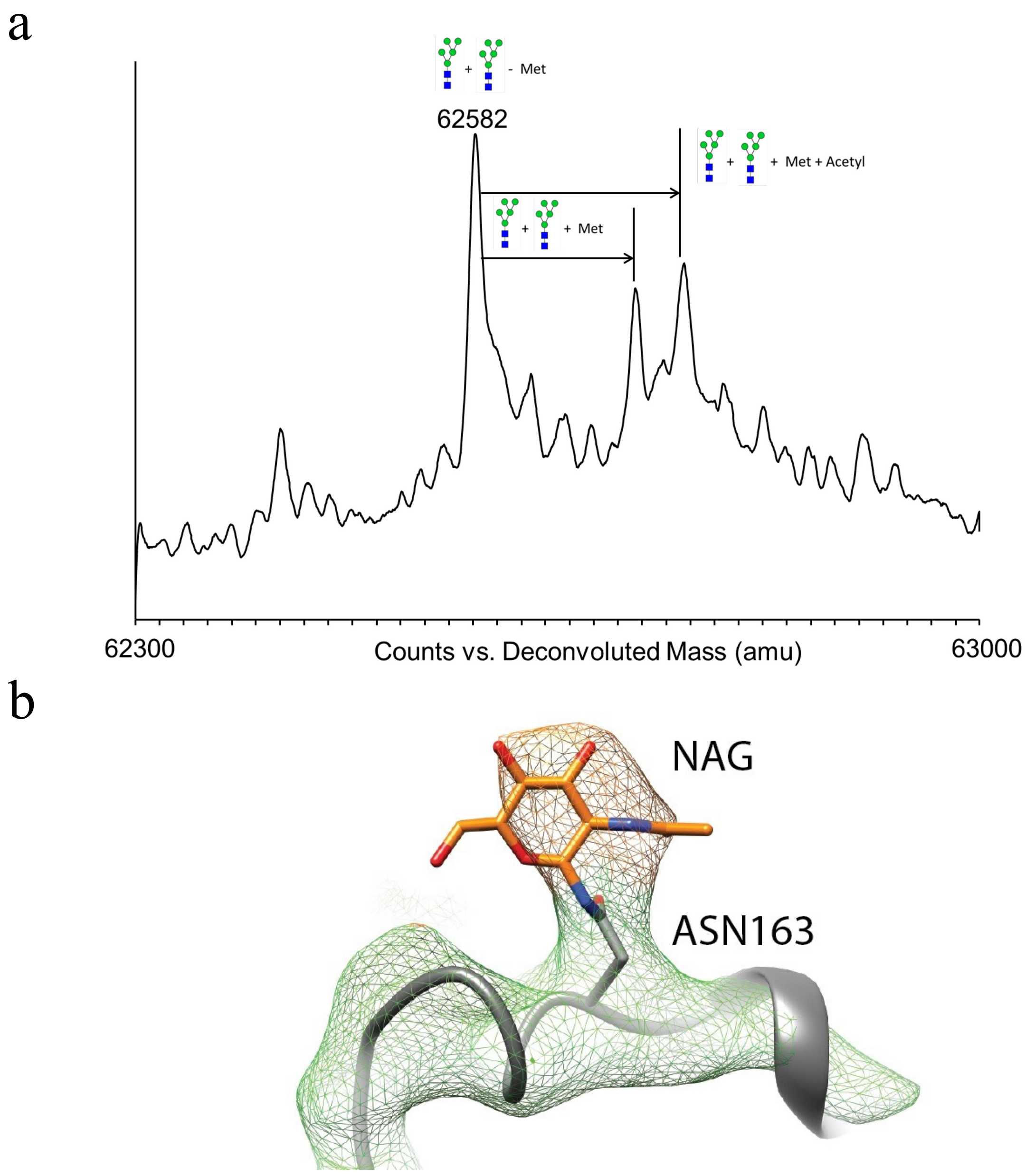

**Figure S8 | Intact Protein Mass Spectrometry. a**, Purified SLC1A5 is analyzed by mass spectrometry. The major observed mass of 62,582 Da is consistent with the addition of two 5 oligo mannose carbohydrate chains and the removal of the N-terminal Met. The other major species observed are 62720 Da (two 5 oligo mannose carbohydrate chains plus the N-term Met) and 62,760 Da (two 5 oligo mannose carbohydrate chains plus the N-term Met plus acetylation). **b**, The electron density (8σ) indicating the N-linked glycosylation on Asn163 is shown in mesh in the context of the atomic model with side chains shown as sticks and the backbone as cartoon. N-acetyl-D-glucosamine linked to Asn163 is colored in orange.

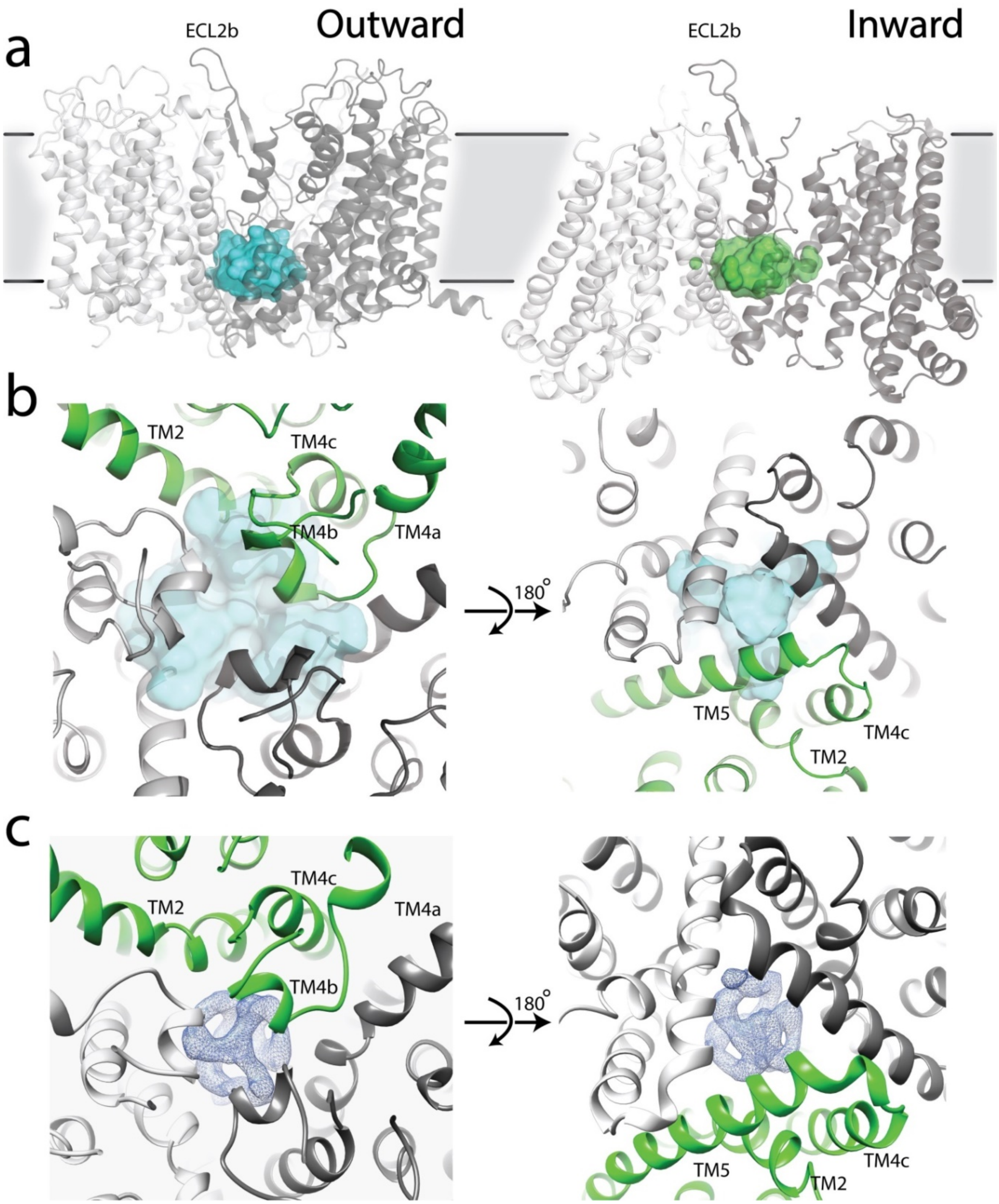

**Figure S9 | Internal cavity of SLC1A5. a**, View from the membrane, one internal cavity is observed from both the outward- (left, shown in in cyan surface) and inward-facing (right, shown in green surface from PDB: 6GCT) states of SLC1A5 cryo-EM structures. **b**, Close-up views of inter-protomer interactions via TM2, TM4 and TM5 in the outward-facing state with the internal cavity shown in cyan surface from the extracellular side (left) and the cytoplasm (right). **c**, None protein-like densities are observed inside the cavity and shown in blue meshes (4σ) from the extracellular side (left) and the cytoplasm (right).

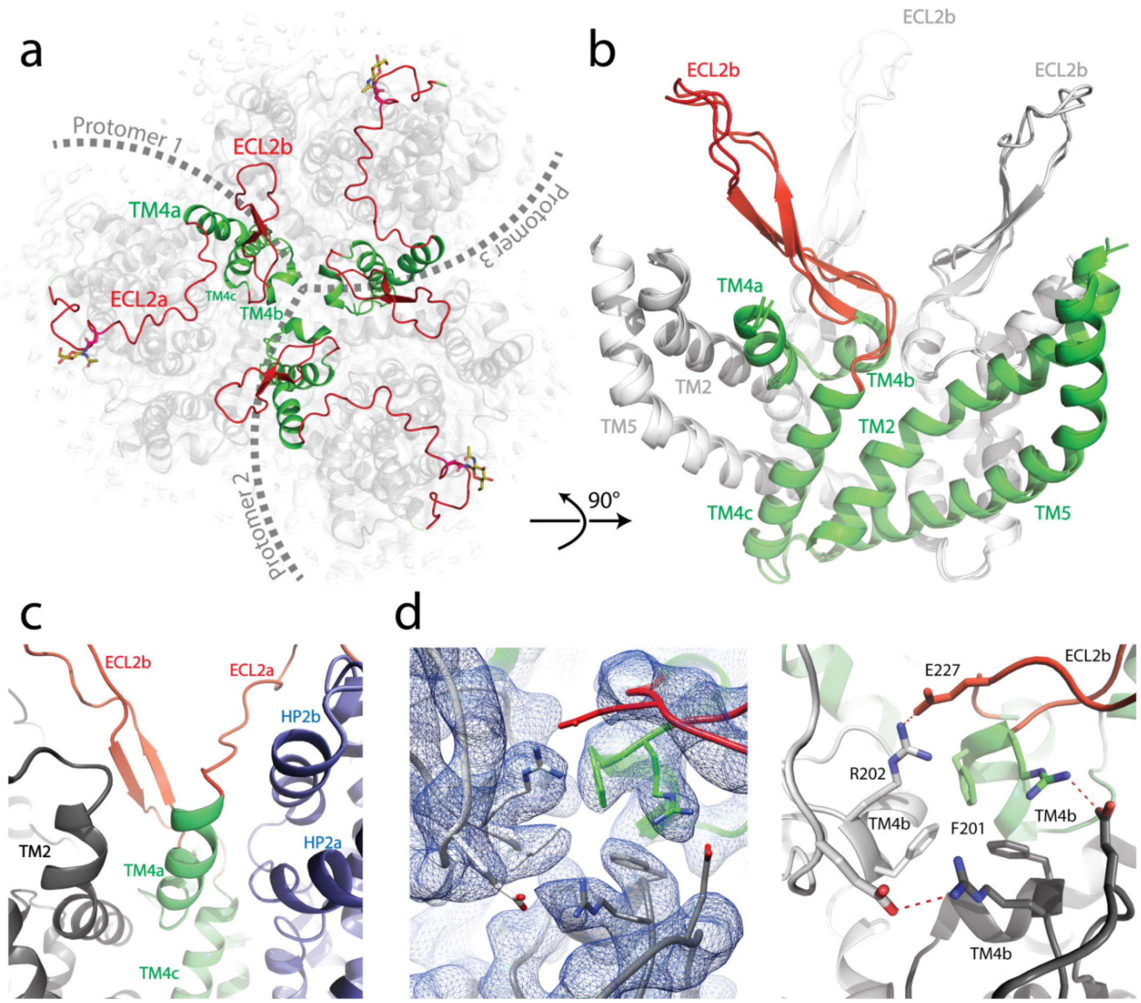

**Figure S10 | Inter-protomer interaction of SLC1A5 a**, View of trimer from the extracellular side of the membrane highlighting the ECL2 (red) and the TM4 (green). Boundaries for each protomer are indicated by gray dashes. TM4 forms the major contact in the center region of SLC1A5 trimer and is composed by three α helices. Three TM4b form the direct interactions in the SLC1A5 trimer. **b**, Superposition of outward-facing state with inward-facing state (PDB:6GCT) of SLC1A5 viewed from membrane showing ECL2b. One beta-turn (ECL2b colored in red) locates between TM4b and TM4c. TM4a-c together with TM2 and TM5 form the center frame in the SLC1A5 trimer. The part of the central core and the ECL2b from one protomer of the SLC1A5 trimer are colored in green and red, respectively, with the rest in gray. **c**, Close-up view of inter-protomer interaction of outward-facing structure viewed from the membrane. TM4a and TM4c interact with the transport domain from the same protomer (colored in slate) and the TM2 from the neighboring protomer (colored in black). **d**, Inter-protomer interaction along 3-fold axis involving residues, Arg202, Glu227 and Phe201 in outward-facing structure. Electron density map was represented in blue mesh (8σ, left). Hydrogen bond interactions are indicated by red dashes (right). Note that the residue Glu shows partial coverage of the electron density on the carboxylic acid group at contour level of 8σ.

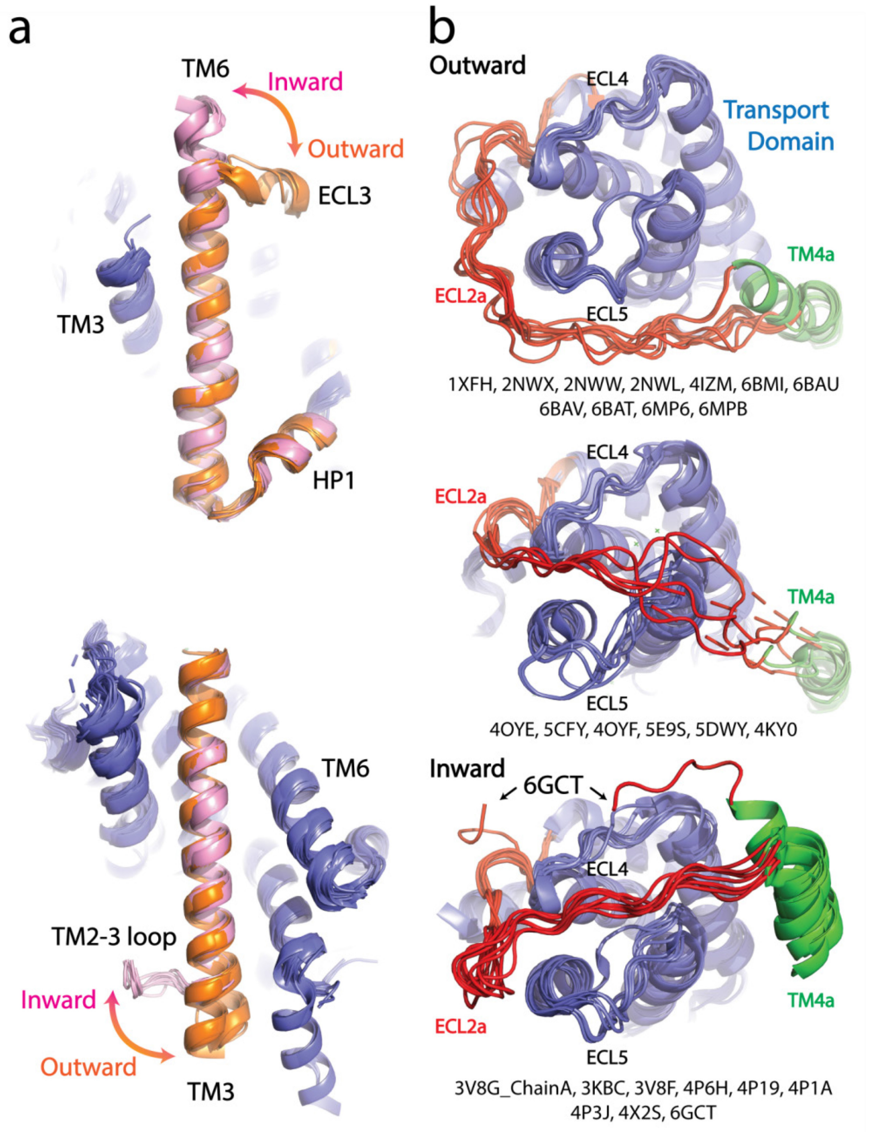

**Figure S11 | Superposition of transport domains from the available structures.** **a**, A close-up view of TM3 and TM6 of transport domains. The transport domains are colored in slate. The TM6, TM3 and part of the HP1a are colored in orange in the outward-facing state and pink in the inward-facing state. Compared to the outward-facing state, the TM6 extends the length by approximately 1.5 α-helical turn from the ECL3 sequence in the inward-facing state (top). In the inward-facing state, the TM3 is shorter by approximately 1.5 α-helix unwinding into the TM2-3 loop compared to the outward-facing state (bottom). **b**, Transport domains viewed from extracellular face. The transport domains and ECL2a are colored in slate and red, respectively. In the outward-facing state, the ECL2a can be situated at the side of the transport domain (top) or crossing over the crests of ECL4 and ECL5 (middle). In the inward-facing state, the ECL2a forms a bent and is repositioned to extend along the ECL4 and ECL5 (bottom). Note that the ECL2a from 3V8G_ChainC, 5MJU, 5LM4, 5LLM, and 5LLU were largely unmodeled and were not included into this analysis. The binding of Fabs or crystal contacts may induce conformational changes on the ECL2a region.

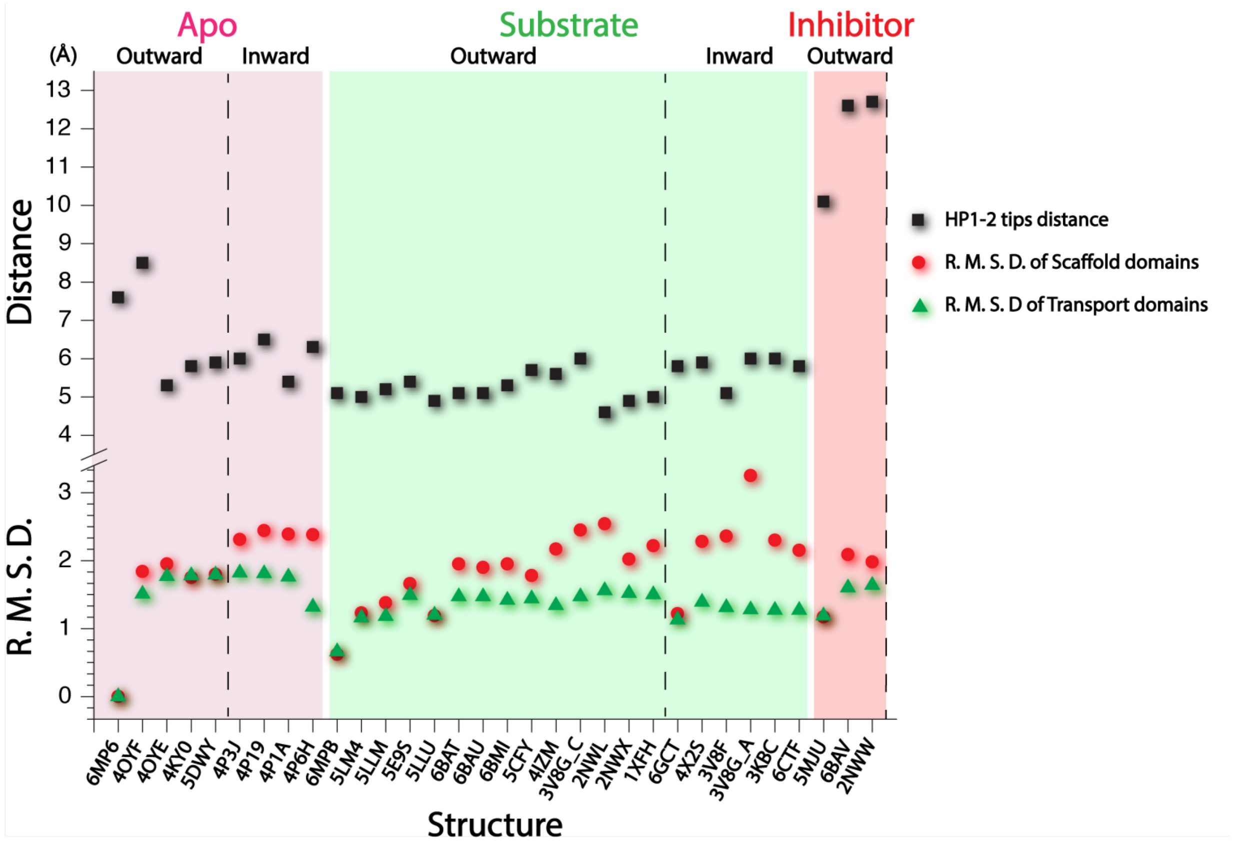

**Figure S12 | The distances of HP1-2 tips and R.M.S.D. of Cα-alignment corresponding to data in Table S2.** All available structures were divided into three groups: apo form in pink; substrate binding form in green; competitive inhibitor binding form in red. The groups are further divided into the outward- and inward-facing states by the dash lines. Black squares are the distances of HP1-2 tips measured using the Cα atoms from S352 and P432 for SLC1A5 structures; S344 and P424 for SLC1A3 structures; S277 and P356 for $Glt_{PH}$ structures; S279 and P359 for $Glt_{TK}$ structures. The R.M.S.D. values were calculated using the secondary-structure matching (SSM). The red circles and green triangles are the R.M.S.D values between SLC1A5-cKM4012 (Fab) and the other structures, calculated using the scaffold and transport domains respectively. Note: structures of 4P19, 4P3J, and 4OYE were determined without substrate and sodium ions in the substrate binding pocket.

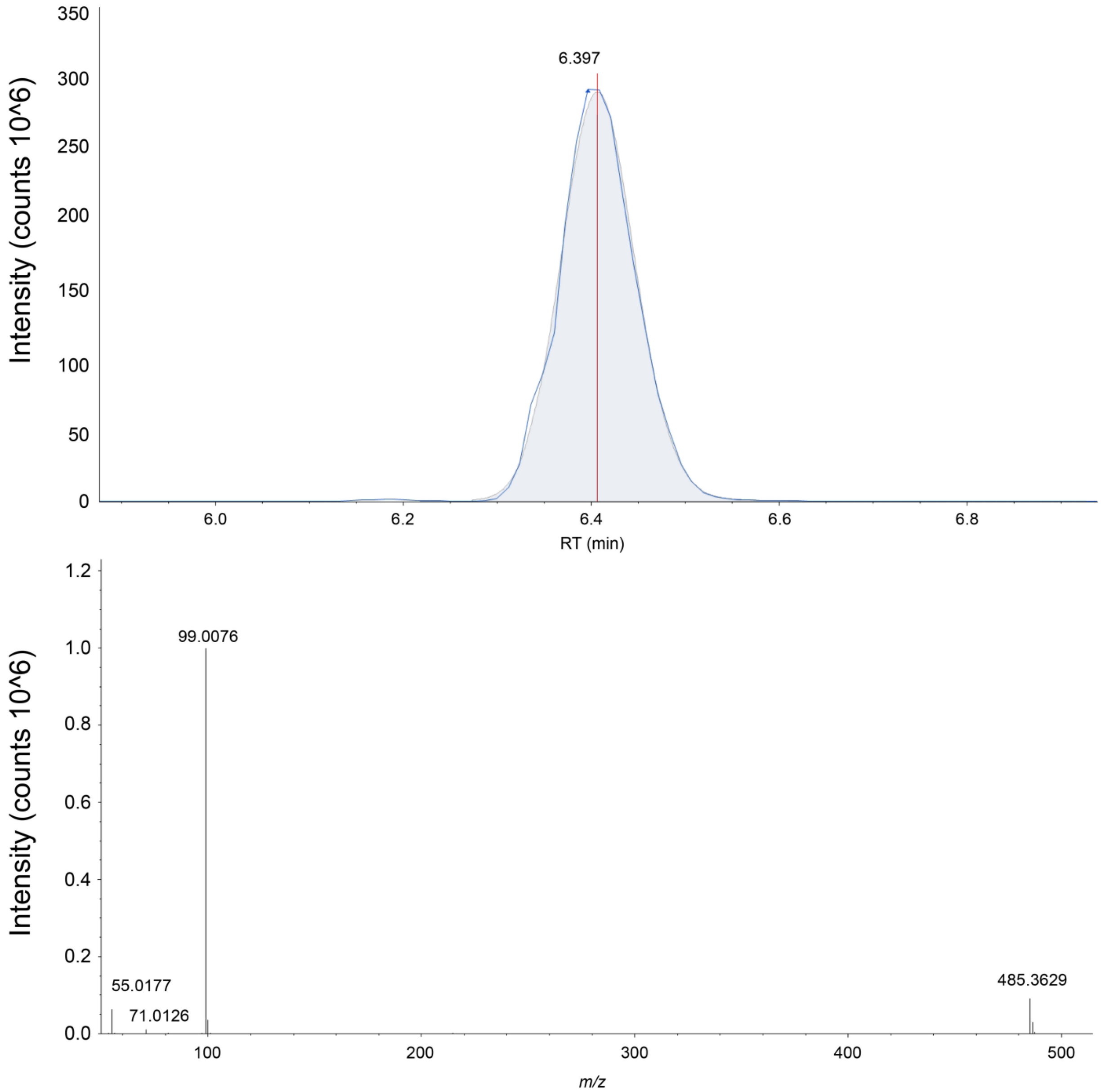

**Figure S13 | The extracted ion chromatograph (above) and the mass spectrum (below) for CHS is shown in negative ion mode**. CHS is measured with a 3 ppm accuracy in the SLC1A5 sample. The exact mass, elution time, and fragmentation pattern are validated against a commercially bought standard (Sigma-Aldrich, St. Louis)**.**

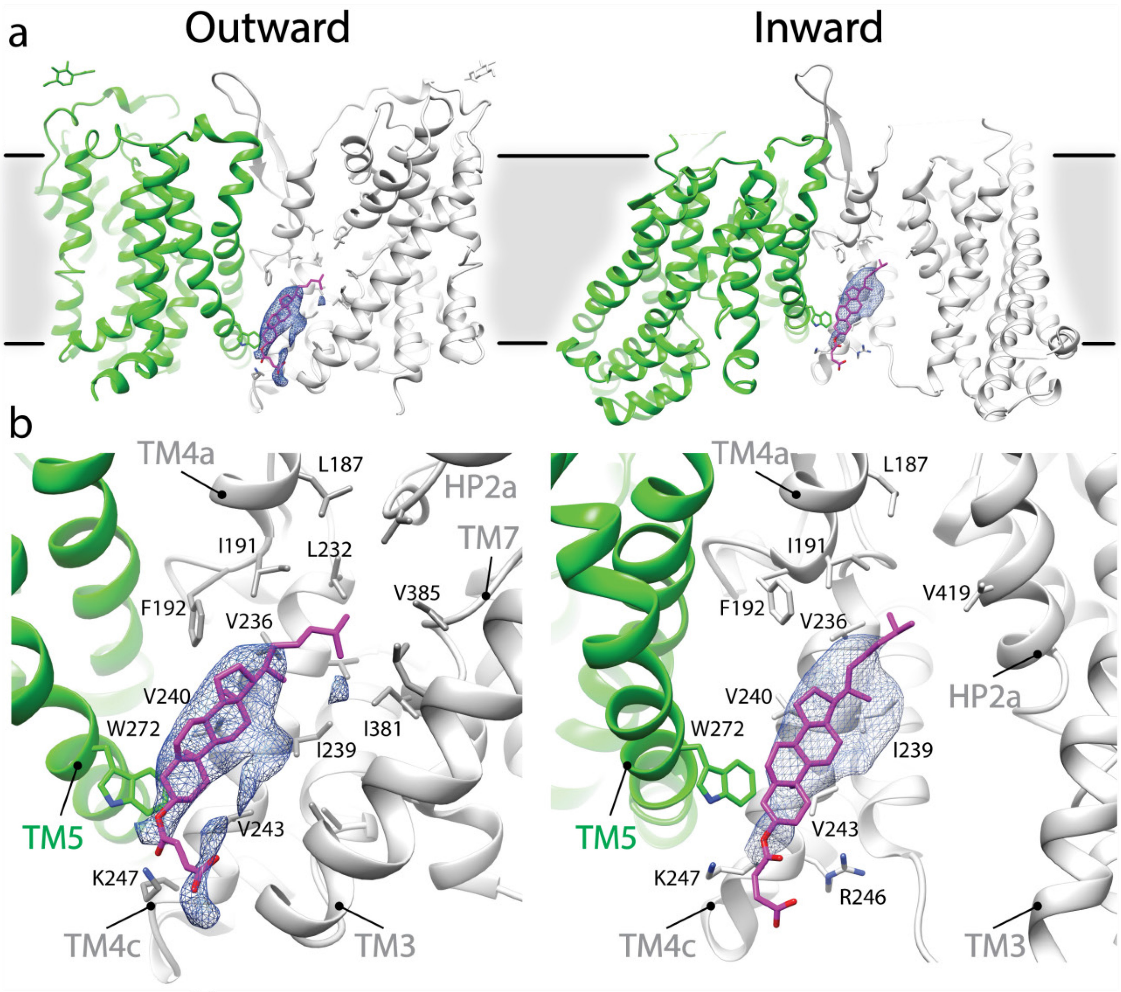

**Figure S14 | Potential CHS interaction in outward- and inward-facing conformation of SLC1A5. a**, Elongated densities, shown as blue mesh (~6σ), are observed between two protomers of SLC1A5 trimer in both the outward- and inward-facing (PDB: 6GCT) cryo-EM structures. **b**, A density is embedded in the cavity enclosed by two SLC1A5 protomers (one in green, and the other in gray). One CHS molecule, shown as magenta sticks, can be docked into this density. TM4a and TM4c provide the primary accommodation site for the CHS molecule. The contour and composition of the cavity define the orientation of the bound CHS. Note that a cholesterol molecule can be accommodated in the hydrophobic pocket the same as CHS.

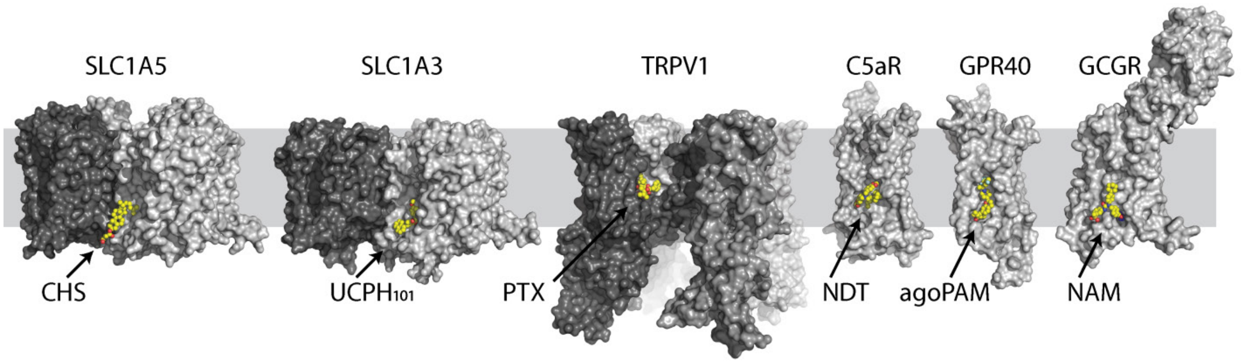

**Figure S15 | Selective examples of allosteric druggable pockets at the lipid-exposed surface near the intracellular part of membrane proteins.** Different protomers are shown in surface with different gray scale. Each ligand is colored in yellow ball-and-sticks. Examples of allosteric sites include the RTX agonist binding site in the capsaicin and heat-activated cation channel TRPV1 (17), the extra-helical NDT9513727 antagonist binding site in the complement C5a receptor (18), the agoPAM binding site in the free fatty acid receptor GPR40 (19) and a negative allosteric modulator binding site in the glucagon receptor (20).

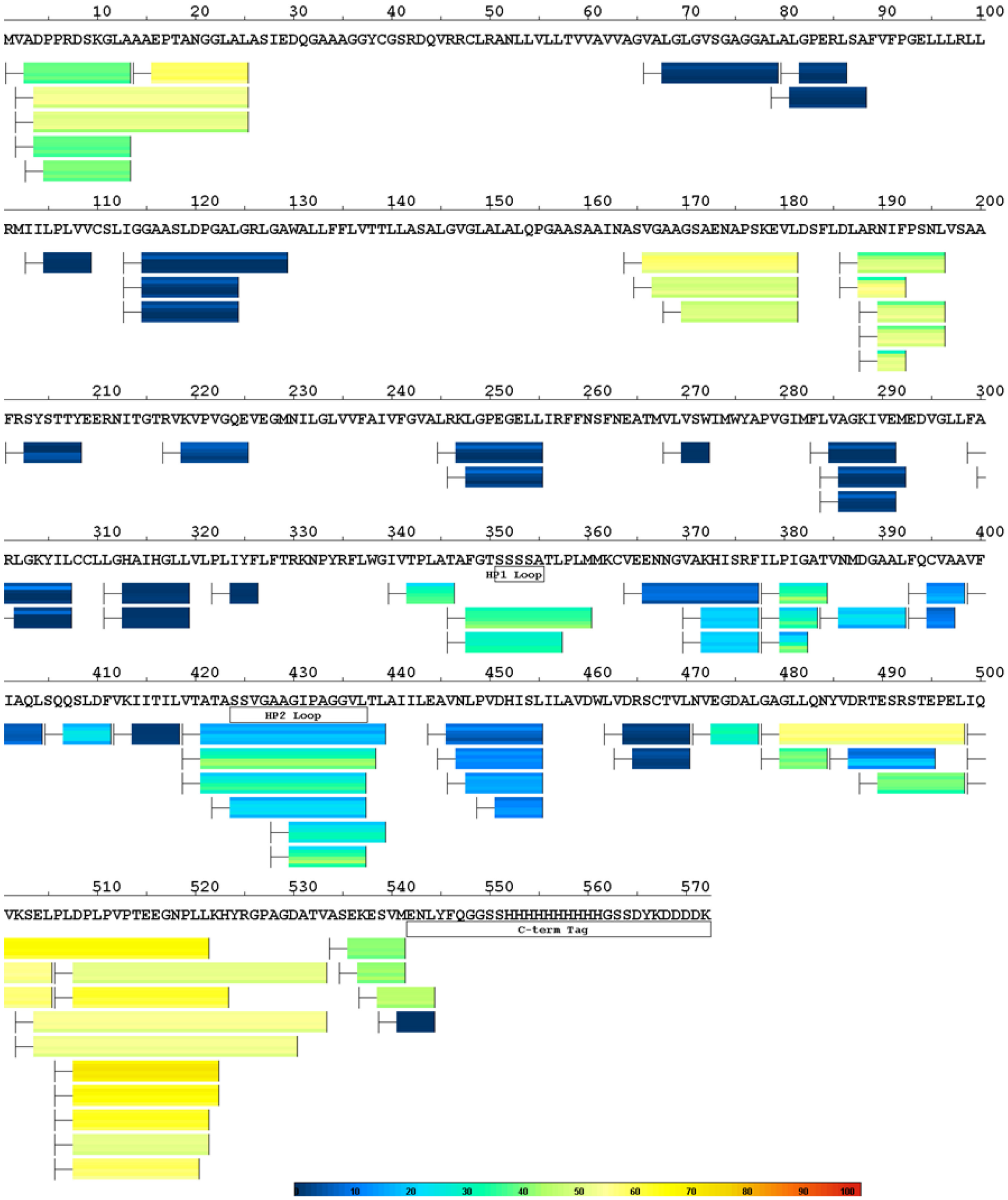

**Figure S16 | SLC1A5 Dynamics**. **a,** HDX heat map of SLC1A5 mapped below its sequence. Peptides analyzed in the HDX experiment are indicated by rectangular boxes below the transporter sequence. Each box is subdivided into six sections representing each of the six exchange time points. The percent deuterium exchange is indicated for each peptide time point according to the colored key. For each peptide, the first 2 N-terminal amino acids were excluded from the analysis due to rapid back-exchange.

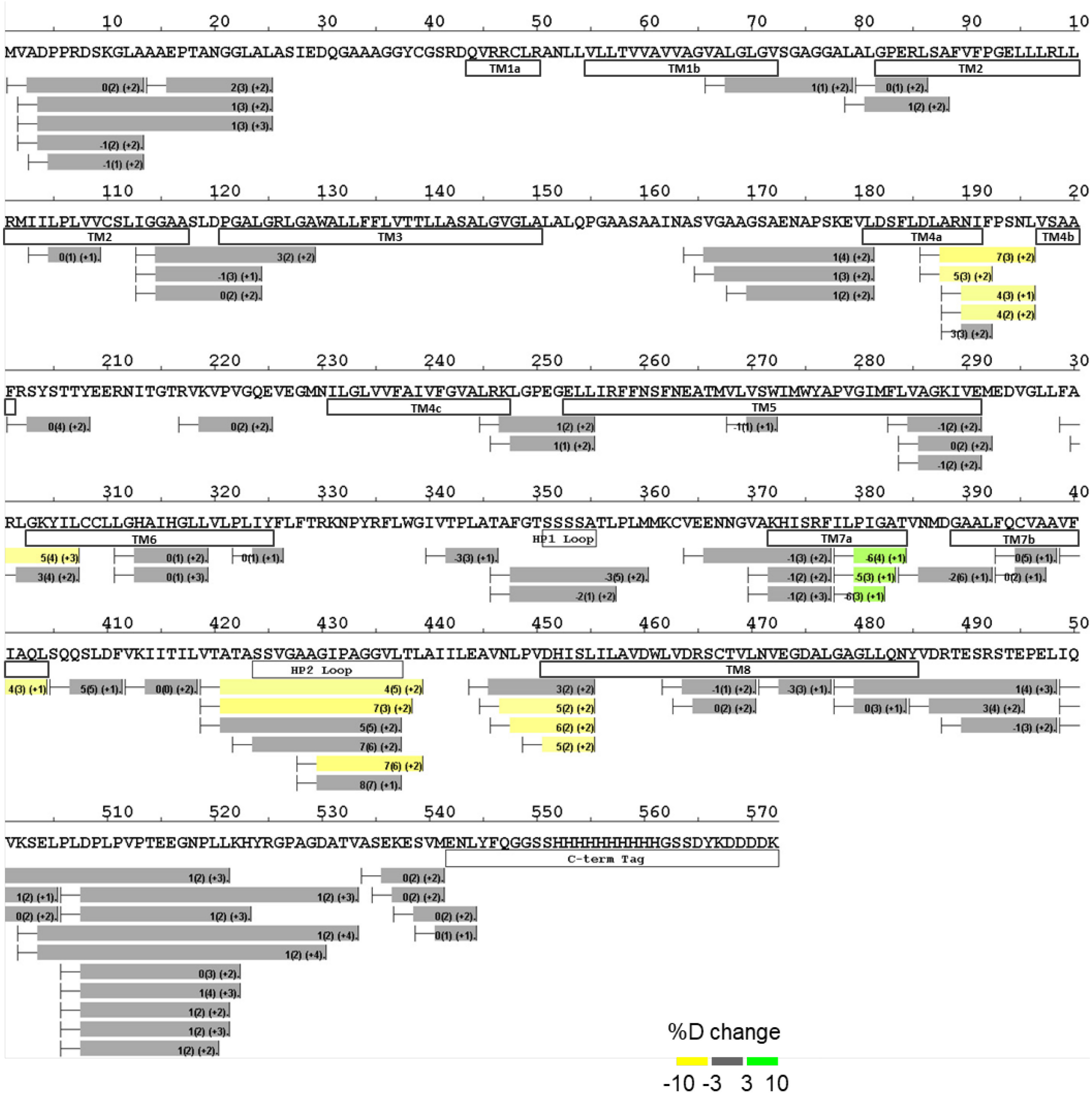

**Figure S16 | SLC1A5 Dynamics**. **b**, HDX perturbation data of gutamine binding mapped to the SLC1A5 sequence. Peptides analyzed in the HDX experiment are indicated by rectangular boxes below the construct sequence. The number in each box represents the change in the average deuterium uptake upon compound binding across all 6 time points. Standard error and peptide ion charge state are also noted in parenthesis. The peptide boxes are colored based on deuterium uptake differences according to the colored key. For each peptide, the first 2 N-terminal amino acids were excluded from the analysis due to rapid back-exchange. Structural features are mapped directly below the sequence. See Figure S11 for a more detailed map of structural features.

**Table S1. Data collection, reconstruction, and model refinement statistics**

| | SLC1A5_cKM4012<br>EMD-9187, PDB: 6MP6 | SLC1A5_cKM4012_L_Gln<br>EMD-9188, PDB: 6MPB |
|---|---|---|
| **Data collection** | | |
| Microscope | Titan Krios | Titan Krios |
| Voltage (keV) | 300 | 300 |
| Nominal magnification | 22,500 x | 22,500 x |
| Exposure navigation | Stage Position | Stage Position |
| Electron exposure (e /Å$^2$) | 42 | 42 |
| Dose rate (e/pixel/sec) | 5 | 5 |
| Detector | K2 Summit | K2 Summit |
| Pixel size (Å)* | 0.543 | 0.543 |
| Defocus range (μm) | 1.2 to 2.5 | 1.2 to 2.5 |
| Micrographs Used | 3804 | 5228 |
| Final Refined particles (no.) | 165,067 | 253,220 |
| **Reconstruction** | | |
| Symmetry imposed | C1 | C1 |
| Resolution (global) | | |
| FSC 0.143 | 3.50 Å | 3.83 Å |
| Applied B-factor (Å$^2$) | -15 | -15 |
| **Refinement** | | |
| Protein residues | 1341<br>(SLC1A5) | 1344<br>(SLC1A5_L_Gln) |
| Map Correlation Coefficient | 0.815 | 0.785 |
| R.m.s deviations | | |
| Bond lengths (Å) | 0.006 | 0.005 |
| Bond angles (°) | 1.085 | 1.062 |
| Ramachandran | | |
| Outliers | 0.00 % | 0.00 % |
| Allowed | 7.88 % | 8.93 % |
| Favored | 92.12 % | 91.07 % |
| Poor rotamers (%) | 0.00 % | 0.00 % |
| MolProbity score | 1.62 | 1.67 |
| EMRinger score | 2.59 | 1.99 |
| Clashscore (all atoms) | 3.44 | 3.67 |

*Calibrated pixel size at the detector

|  | Name | PDB Code | Organism | Method | Resolution (Å) | Conformation | HP1-2 tip distance | Ligand | Sequence Identity (%) | Cα R.S.M.D. (Å) Global | Scaffold | Transport |
|---|---|---|---|---|---|---|---|---|---|---|---|---|
| 1 | SLC1A5 | 6MP6 | Homo sapiens | Cryo EM | 3.5 | Outward | 7.6 | Fab | 100.0 | 0.00 | 0.00 | 0.00 |
| 2 | SLC1A5 | 6MPB | Homo sapiens | Cryo EM | 3.8 | Outward | 5.1 | Fab, Gln | 100.0 | 0.65 | 0.62 | 0.66 |
| 3 | SLC1A5 | 6GCT | Homo sapiens | Cryo EM | 3.9 | Inward | 5.8 | Gln | 100.0 | N/A | 1.22 | 1.13 |
| 4 | SLC1A3 | 5LM4 | Homo sapiens | X ray | 3.1 | Outward | 5.0 | UCPH101, Asp | 56.5 | 1.26 | 1.23 | 1.16 |
| 5 | SLC1A3 | 5LLM | Homo sapiens | X ray | 3.3 | Outward | 5.2 | UCPH101, Asp | 56.4 | 1.29 | 1.38 | 1.18 |
| 6 | SLC1A3 | 5LLU | Homo sapiens | X ray | 3.3 | Outward | 4.9 | Asp | 56.5 | 1.38 | 1.19 | 1.20 |
| 7 | SLC1A3 | 5MJU | Homo sapiens | X ray | 3.7 | Outward | 10.1 | TFB-TBOA, UCPH101 | 56.4 | 1.23 | 1.17 | 1.19 |
| 8 | GltPh | 6BAT | Pyrococcus horikoshii | X ray | 3.4 | Outward | 5.1 | Asp | 25.9 | 2.13 | 1.95 | 1.47 |
| 9 | GltPh | 6BAV | Pyrococcus horikoshii | X ray | 3.7 | Outward | 12.6 | Benzylcystein | 26.0 | 2.18 | 2.09 | 1.60 |
| 10 | GltPh | 6BAU | Pyrococcus horikoshii | X ray | 3.8 | Outward | 5.1 | Cys | 26.0 | 2.12 | 1.90 | 1.47 |
| 11 | GltPh | 6BMI | Pyrococcus horikoshii | X ray | 3.9 | Outward | 5.3 | Ser | 26.0 | 2.08 | 1.95 | 1.42 |
| 12 | GltTk | 5E9S | Thermococcus kodakarensis | X ray | 2.8 | Outward | 5.4 | Asp | 30.5 | 2.08 | 1.66 | 1.49 |
| 13 | GltTk | 5DWY | Thermococcus kodakarensis | X ray | 2.7 | Outward | 5.9 | Apo | 30.5 | 2.31 | 1.80 | 1.79 |
| 14 | GltPh | 4X2S | Pyrococcus horikoshii | X ray | 4.2 | Inward | 5.9 | Asp | 26.4 | N/A | 2.28 | 1.39 |
| 15 | GltPh | 4OYF | Pyrococcus horikoshii | X ray | 3.4 | Outward | 8.5 | Apo | 25.9 | 2.03 | 1.84 | 1.51 |
| 16 | GltPh | 5CFY | Pyrococcus horikoshii | X ray | 3.5 | Outward | 5.7 | Asp | 26.2 | 2.23 | 1.78 | 1.44 |
| 17 | GltPh | 4P3J | Pyrococcus horikoshii | X ray | 3.5 | Inward | 6.0 | Apo | 25.9 | N/A | 2.31 | 1.82 |
| 18 | GltPh | 4P1A | Pyrococcus horikoshii | X ray | 3.8 | Inward | 5.4 | TL, HG | 25.9 | N/A | 2.39 | 1.76 |
| 19 | GltPh | 4P19 | Pyrococcus horikoshii | X ray | 3.8 | Inward | 6.5 | Apo, HG | 25.9 | N/A | 2.44 | 1.81 |
| 20 | GltPh | 4OYE | Pyrococcus horikoshii | X ray | 4.0 | Outward | 5.3 | Apo | 25.3 | 2.31 | 1.95 | 1.77 |
| 21 | GltPh | 4P6H | Pyrococcus horikoshii | X ray | 4.1 | Inward | 6.3 | TL, HG | 25.9 | N/A | 2.38 | 1.32 |
| 22 | GltTk | 4KY0 | Thermococcus kodakarensis | Xray | 3.0 | Outward | 5.8 | Apo | 30.5 | 2.21 | 1.75 | 1.78 |
| 23 | GltPh | 4IZM | Pyrococcus horikoshii | X ray | 4.5 | Outward | 5.6 | Asp, HG | 25.9 | 2.28 | 2.17 | 1.34 |
| 24 | GltPh | 3V8F | Pyrococcus horikoshii | X ray | 3.8 | Inward | 5.1 | Asp, HG | 26.0 | N/A | 2.36 | 1.31 |
| 25 | GltPh | 3V8G, Chain A | Pyrococcus horikoshii | X ray | 4.7 | Inward | 6.0 | Asp | 25.9 | N/A | 3.25 | 1.28 |
| 25 | GltPh | 3V8G, Chain C | Pyrococcus horikoshii | X ray | 4.7 | Outward | 6.0 | Asp | 25.9 | 2.73 | 2.45 | 1.47 |
| 26 | GltPh | 3KBC | Pyrococcus horikoshii | X ray | 3.5 | Inward | 6.0 | Asp, HG | 26.2 | N/A | 2.30 | 1.27 |
| 27 | GltPh | 2NWL | Pyrococcus horikoshii | X ray | 3.0 | Outward | 4.6 | Asp | 25.9 | 2.05 | 2.54 | 1.56 |
| 28 | GltPh | 2NWW | Pyrococcus horikoshii | X ray | 3.2 | Outward | 12.7 | TBOA | 25.9 | 2.34 | 1.98 | 1.64 |
| 29 | GltPh | 2NWX | Pyrococcus horikoshii | X ray | 3.3 | Outward | 4.9 | Asp | 25.9 | 2.00 | 2.02 | 1.52 |
| 30 | GltPh | 1XFH | Pyrococcus horikoshii | X ray | 3.5 | Outward | 5.0 | Asp | 25.9 | 2.23 | 2.22 | 1.50 |
| 31 | GltPh | 6CTF | Pyrococcus horikoshii | Xray | 4.1 | Inward | 5.8 | Asp | 25.9 | N/A | 2.15 | 1.27 |

**Table S2. Overview of the available structures of human SLC1 family and prokaryotic homologues.** The structures were ranked by the release date except for 6CTF. HP1-2 tip distance is measured using the Cα atoms from S352 and P432 for SLC1A5 structures; S344 and P424 for SLC1A3 structures; S277 and P356 for GltPh structures; S279 and P359 for GltTk structures. Sequence identity is calculated using the LALIGN program with the protein sequence of human SLC1A5 as the reference. Cα R.M.S.D. is calculated using the secondary-structure matching (SSM). The global, scaffold, and transport domains structures of SLC1A5-cKM4012 (Fab) serve as the references, respectively. Note: structures of 4P19, 4P3J, and 4OYE were determined without substrate and sodium ions in the substrate binding pocket.
